# The genomic and cultural diversity of the Inka Qhapaq hucha ceremony in Chile and Argentina

**DOI:** 10.1101/2024.02.19.581063

**Authors:** Constanza de la Fuente Castro, Constanza Cortés, Maanasa Raghavan, Daniela Castillo, Mario Castro, Ricardo Verdugo, Mauricio Moraga

**Affiliations:** Departamento de Genética Humana, ICBM, Universidad de Chile, Av Independencia 1027, Santiago, 8380453, Región Metropolitana, Chile.; Department of Human Genetics, University of Chicago, 920 E 58th St, Chicago, 60637, IL, USA.; Escuela de Arqueología, Universidad Austral, Liborio Guerrero 1765, Puerto Montt, 5504327, Región de Los Lagos, Chile.; Museo Nacional de Historia Natural, Parque Quinta Normal, Santiago, 8500000, Región Metropologitana, Chile.; Departamento de Morfología, Facultad de Medicina, Clínica Alemana-Universidad del Desarrollo, Av. Plaza 680, Santiago, 7610615, Región Metropolitana, Chile.; Departamento de Oncología Básico-Clínica, Universidad de Chile, Av. Independencia 1027, Santiago, 8380453, Región Metropolitana, Chile.; Instituto de Investigación Interdisciplinaria, Universidad de Talca, 2 Norte 685, Talca, 3465548, Región del Maule, Chile.; Departamento de Antropología, Facultad de Ciencias Sociales, Universidad de Chile, Av. Capitán Ignacio Carrera Pinto 1045, Santiago, 7800284, Región Metropolitana, Chile.

**Keywords:** Paleogenomics, Andes, Inka, Population genomics

## Abstract

The South American archaeological record has ample evidence of the socio-cultural dynamism of human populations in the past. This has also been supported through the analysis of ancient genomes, by showing evidence of gene flow across the region. While the extent of these signals is yet to be tested, the growing number of ancient genomes allows for more fine-scaled hypotheses to be evaluated. In this study, we assessed the genetic diversity of individuals associated with the Inka ritual, Qhapaq hucha. As part of this ceremony, one or more individuals were buried with Inka and local-style offerings on mountain summits along the Andes, leaving a very distinctive record. Using paleogenomic tools, we analyzed three individuals: two newly-generated genomes from El Plomo Mountain (Chile) and El Toro Mountain (Argentina), and a previously published genome from Argentina (Aconcagua Mountain). Our results reveal a complex demographic scenario with each of the individuals showing different genetic affinities. Furthermore, while two individuals showed genetic similarities with present-day and ancient populations from the southern region of the Inka empire, the third individual may have undertaken long-distance movement. The genetic diversity we observed between individuals from similar cultural contexts supports the highly diverse strategies the Inka implemented while incorporating new territories. More broadly, this research contributes to our growing understanding of the population dynamics in the Andes by discussing the implications and temporality of population movements in the region.

## Introduction

The South American archaeological record has ample evidence of the profound socio-cultural and demographic dynamism of past human populations in the region. This dynamism intensified in later periods, particularly with the establishment of agricultural and ceramic technologies [1]. From a genetic perspective, mitochondrial DNA (mtDNA) and genome-wide data have suggested different degrees of population continuity [e.g. 2,3] as well as evidence of gene flow and population interactions at both broad and regional scales. Examples of broad-scaled interactions include: i) genetic affinity between ancient individuals from the Channel Islands in California and the Central Andes by 4,200 BP [4]; ii) evidence of additional Mesoamerican-related ancestry in several South American populations [5]; and iii) a north-to-south gradient of ancestry associated with populations from central Chile in South Patagonian populations [6]. Meanwhile, most of the interactions at a regional scale have been associated with empire states such as Tiwanaku and Inka. In the case of Tiwanaku, this has been reflected as an excess of allele sharing between ancient individuals from Tiwanaku administrative center and the highland region of South Peru compared to other regions [2]. In addition, ancient genomes from Cusco analyzed by Nakatsuka et al. (2021) and dated to the Inka period, showed genetic heterogeneity, reflecting the cosmopolitan nature of the empire’s capital [2]. Similarly, a recent paleogenomics study of the Machu Picchu site showed that individuals buried there “exhibited ancestries from throughout the Inca Empire” [7].

Of all the cultural developments in South America, the Inka stands out as one of the most monumental and largest empires in the region. Expanding from its capital Cusco around the 14th century CE, the Inka empire, or *Tawantinsuyu*, rapidly spread north, reaching present-day southern Colombia, and south, reaching the central-south region of present-day Chile and central-west Argentina [8,9]. Evidence of its advance is well documented through different elements, especially styles, and iconography on different materials such as ceramics and textiles. In addition, the *Qhapaq ñam*, or the Inka trail, is an extensive network of roadways that bears witness to the high connectivity and expanded influence of the Inka throughout the Andes [10].

The *Tawantinsuyu* encompassed a territory of nearly 1,000,000 km² and was divided into four major regions: *Chinchaysuyu* (north), *Antisuyu* (east), *Collasuyu* (south), and *Cuntisuyu* (west), exhibiting significant variability in regional human populations, landscapes, and climates. This diversity not only provided the *Tawantinsuyu* access to a wide range of resources, but also necessitated interactions with diverse populations with distinct political, social, cultural, and economic practices. As a result, the Inka devised varying annexation strategies for each region [11], as evidenced in the archaeological record. The dynamics of interaction and domination fluctuated, based on both the *Tawantinsuyu*’s interests and the unique characteristics and motivations of the local populations [11–13]. Moreover, there is evidence of simultaneous implementation of different coordinated policies to assert dominance over a single territory [e.g. 10–12].

During the Inka expansion, some of the highest mountains in the Andes became places of special meaning, reaching their highest expression through a ritual known as *Qhapaq hucha*. As part of this ritual, one or more individuals, usually younger than 16 years old, were buried close to the summit, together with an assortment of grave goods of local and foreign origin. It has been proposed that these sacrifices marked the culmination of a ceremonial pilgrimage that originated in the heart of the *Tawantinsuyu* capital, Cusco [17]. Across the empire’s range, there are at least 14 summits with human burials and the region of *Collasuyu* stands out for its high incidence. Accompanying these human offerings are secondary tributes such as camelid or anthropomorphic figurines crafted from *Spondylus* shells, minerals such as silver or gold alloys, food items, coca leaves, feathers, textiles, and pottery [17–21].

The presence of high-altitude shrines has often been interpreted as being indicative of Inka influence and dominion. Through rituals conducted at the base and summit of mountains, the Inka not only appropriated these spaces but also reshaped the ritual landscape, forging stronger connections with local populations [19,22,23]. Interpretations of the implementation and motives for conducting *Qhapaq hucha* across the Inka territory vary widely. These rituals have been seen as demonstrations of dominion over newly acquired lands, as acts of foundational importance, and as preventative measures against disasters like earthquakes, volcanic eruptions, and droughts. Additionally, they have been associated with specific events such as fertility ceremonies for livestock and crops [17]. The presence of these rituals has also been correlated with areas rich in mineral resources [24] or with regions where Inka influence had recently been established, resulting in limited administrative centers and structures dedicated to solar ritual ceremonies [25]. Importantly, little is known about the individuals buried as part of this ceremony, including their origins.

This work aims to investigate the practice and diversity of the *Qhapaq hucha* ceremony through a genetic lens, by characterizing the genetic variation and regional affinities of the individuals associated with these burials. In particular, by implementing a paleogenomic approach, we evaluate the genetic relationships of the individuals associated with the *Qhapaq hucha* ceremony with each other, as well as with other past and present-day populations across South America. More broadly, this research contributes to the understanding of the ancient population dynamics in the Andes by discussing the implications and temporality of population movements in the region.

## Results

Whole-genome sequencing data from three individuals who have been culturally associated with the *Qhapaq hucha* (QH) ceremony in Chile and Argentina were analyzed, of which two were newly sequenced in this study and one was published previously (Table 1).

**Table 1.** Summary of the *Qhapaq hucha* -associated individuals analyzed in this study. See extended sequencing and mapping statistics in Table S1.

| Summit | DoC<sup>1</sup> | Chromosomal sex | mtDNA haplogroup | Y-Chromosome haplogroup |
| --- | --- | --- | --- | --- |
| Cerro El Toro | 3.6x | XY | D1j1a1 | Q1a2a1a1 |
| Aconcagua<sup>2</sup> | 2.4x | XY | C1bi | Q1a2a1a1 |
| Cerro El Plomo | 2.2x | XY | C1b | Q1a2a1a1 |

<sup>1</sup>Depth of coverage
<sup>2</sup>[5]

First, the uniparental signatures of these individuals were analyzed and compared to present-day distributions of the assigned haplogroups in South America. All mitochondrial lineages belonged to known haplogroups in South America (C1b and D1j), some of which show a more restrictive distribution. The lineage C1bi in the Aconcagua individual was first characterized by Gómez-Carballa et al (2015). While this specific lineage was not found in any present-day population, C1b is one of the most frequent lineages in South America [26]. The lineage of the El Plomo individual is also part of C1b but it presents unique variations not overlapping with the ones described for the Aconcagua child. Meanwhile, the lineage D1j1a1 characterized in the El Toro individual shows higher frequencies in northwestern and central Argentina [27–29]. The Y-chromosome haplogroups were characterized for the three individuals, all belonging to the main lineage in the Americas Q1a2a1a1.

The genomes of the three QH individuals were analyzed together with 650 individuals from different present-day populations from Peru, Chile, and Argentina (Figure 1A, Table S2). A principal component analysis (PCA) was performed using smartpca in Eigensoft [30]. The first two principal components (PCs) described two major genetic ancestry gradients in South America: PC1 showing a north-to-south distribution with individuals from Peru at one end and individuals from central-south Chile at the other; and PC2, separating individuals from northern and southern Peru (Figure 1B). The four ancient QH genomes and nearly 200 ancient individuals from Peru, Bolivia, and Chile (see details in Table S3) were projected onto the PC space described in Figure 1B using the lsqproject option in smartpca (Figure 1C). The QH individuals fall at different positions along the genetic gradients, suggesting they have different genetic affinities within South America (Figures 1B and D). Other ancient genomes from South America generally cluster based on their geographical locations, as described elsewhere [2].

**Fig. 1.**
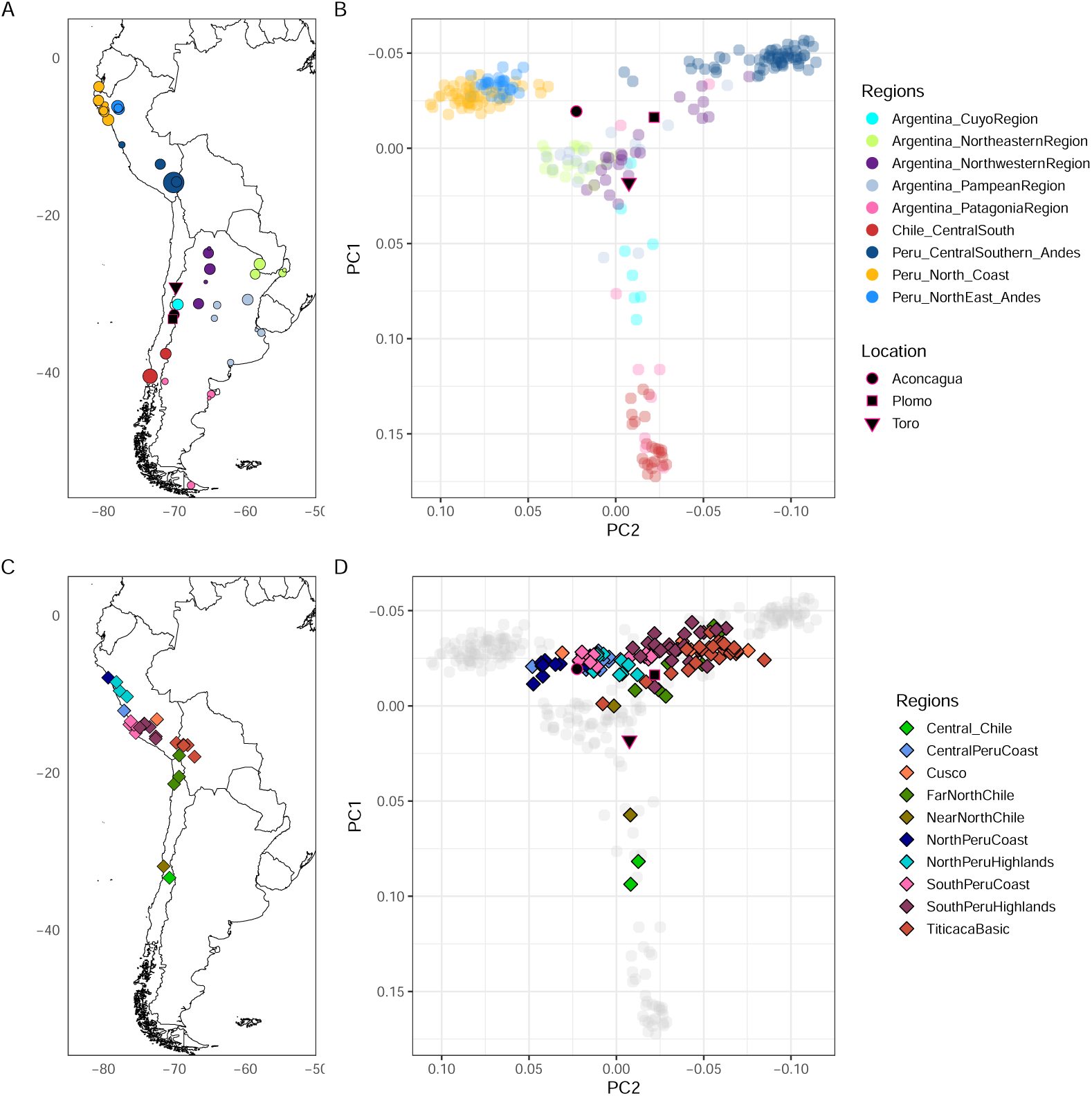
Map and principal component analysis (PCA). A) Locations of the QH-associated burials analyzed in this study. Additionally shown are the geographic distribution of present-day populations included in the analysis, with the size of the circle is indicative of the sample size (the smallest circle is one individual); B) PCA of QH individuals, projected onto PCs 1 and 2 estimated with present-day populations that are color-coded by region; C) Geographic distribution of published ancient genomes included in the analyses; D) PCA of the published ancient genomes and the QH individuals projected onto PCs 1 and 2 estimated with present-day populations.

We evaluated the genetic affinities of the QH genomes to populations from South America using an outgroup-*f* 3 statistic of the form (QH, X; Outgroup), with X representing diverse present-day and ancient individuals from South America. Figure 2 shows the results for only present-day populations as X (see also Figure S1). El Plomo individual shows a stronger association with populations from central-south Peru, while El Toro individual displayed the highest outgroup-*f* 3 values with present-day populations from northwestern and Cuyo region in Argentina and southern latitudes, and Aconcagua shared the greatest genetic affinities with present-day northern and central Coast populations from Peru. The latter has been reported in other studies [2,5].

**Fig. 2.**
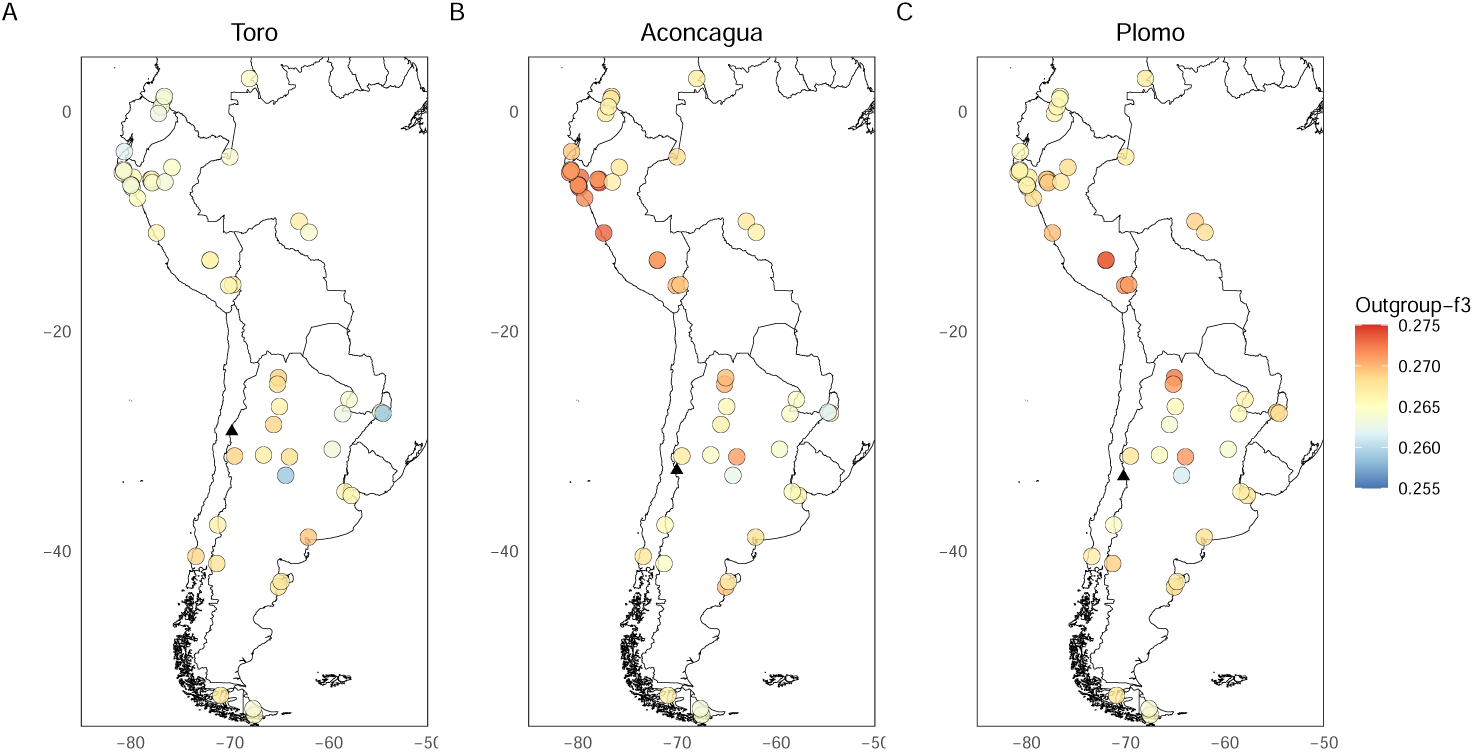
Outgroup-*f* 3 analysis. The analysis was performed with the software admixtools [31] of the form ƒ3(QH, X; Outgroup), where QH represents each of the four ancient genomes associated with the Qhapaq hucha ceremony and X represents different present-day populations from South America. The black triangle shows the geographical location of each QH individual.

In addition, we generated a multi-dimensional scaling (MDS) plot of a distance matrix, based on the outgroup-*f* 3 statistic converted to distance (1 – outgroup-*f* 3), between pairs of ancient individuals from South America, keeping only individuals with at least 10% of the data and the pairs with more than 5,000 overlapping positions (Figure S2-S3). As shown previously [2], we observed that the clustering of ancient individuals mostly followed a pattern based on major geographical regions (e.g. Titicaca basin, south Peru highlands, North Peru highlands). We observed QH individuals clustering with different ancient groups, reflective of the analysis with present-day populations. Overall, the outgroup-*f* 3 analysis suggests that each QH individual shares genetic similarities with different present-day and ancient groups across South America.

To evaluate in detail the different genetic affinities between QH individuals, we implemented a D-statistic analysis of the form D(QH1, QH2; X; Outgroup), where QH1 and QH2 represent pairs of QH individuals and X is a subset of present-day individuals from South America, representing the different regions and genetic clusters on the PCA (Figure 3, Figure S4). Compared to the other QH individuals, the Aconcagua individual is closer to present-day individuals from the north coast of Peru (e.g. Eten). The El Plomo individual mostly shows significant (*|Z| >* 3) allele sharing with present-day individuals from central-south Peru when compared to the El Toro individual. Meanwhile, there is no tested present-day population that is closer to El Toro than to any of the other QH individuals. Similarly, we evaluated the genetic affinities of QH individuals with each other and with other ancient individuals available in the literature using the D-statistic, observing a similar pattern. However, most of the results were not statistically significant with absolute Z-scores lower than 3 (see Figure S5). In contrast to the tests with present-day individuals, we observed two ancient groups that were consistently closer to the El Toro individual: LosRieles 5100BP and Conchalí 700BP. In addition, we re-analyzed shotgun sequencing data generated for three ancient individuals from the same province in Argentina (San Juan) and dated to ca. 1500-2000 cal BP [32]. Despite the low depth of coverage (0.006x to 0.3x) and quality of these genomes, we found their genetic affinities to be similar to the ones observed in the El Toro individual (Figure S6 and S7). Furthermore, while the absolute Z-scores are not statistically significant, they displayed a trend toward the El Toro individual (Figure S5).

**Fig. 3.**
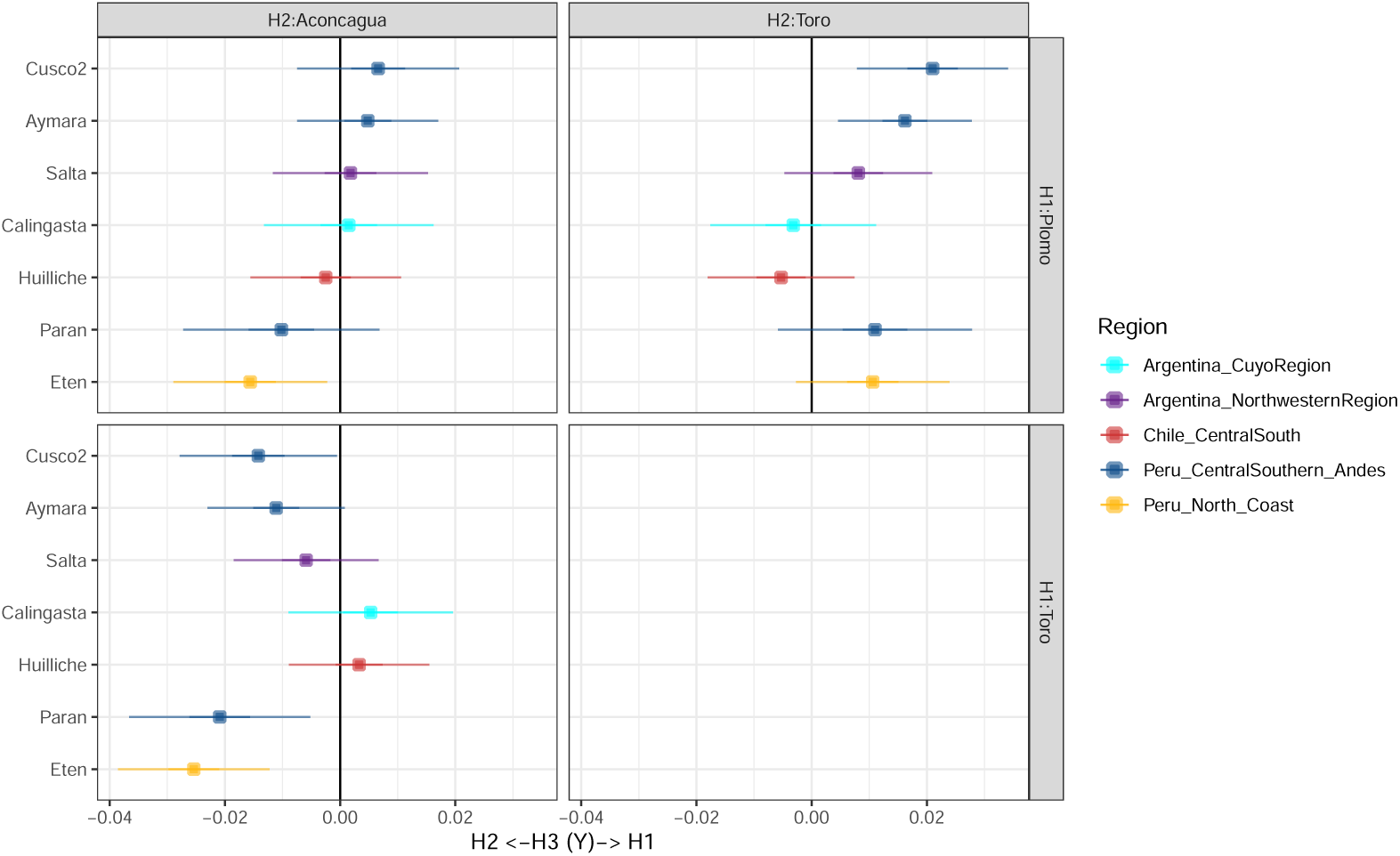
D-statistic of the form D(Qh1, Qh2; X; Outgroup). QH1 and QH2 represent pairs of Qhapaq hucha individuals and X represents different present-day populations from South America. The error bars represent 3 SEs. The analysis was performed using the software admixtools [31]. Extended version in Figure S4.

Using qpWave, we assessed whether QH individuals form a clade with respect to a set of populations referred to as “Right populations” [33] (Table S4). Using a broad geographical and temporal set of right populations (Set1) similar to the model implemented by Posth et al. (2018), all QH individuals form a clade with each other (Figure S8A, p-value *>* 0.05). However, when selecting a more informative list of populations to provide greater regional resolution (Set2), we observed that none of the QH individuals form a clade with each other (Figure S8B, p-value *<* 0.05), supporting the ancestry differences between these individuals reported in the previous analyses. We next used a subset of Set2 with more regionally relevant groups and, again, found that this set was able to distinguish between two pairs of QH individuals, to the exclusion of the El Plomo and El Toro individuals (Set 3 1, Figure S8C). Since LosRieles 5100BP or Conchalí 700BP were the two ancient genomes that were significantly closer to El Toro compared to the other QH individuals in the D-statistics analysis (Figure S5), we reintroduced these two genomes consecutively into Set3 1 to test if we were able to achieve better resolution of the genetic lineage contributing to El Toro (Sets3 2 and 3 3). As expected, we observed that we were only able to differentiate between the El Toro and El Plomo individuals when either the LosRieles 5100BP or Conchalí 700BP genomes were included in the set (Figure S8C-E). This may suggest that the genomes of these two individuals (Los Rieles dated 5,100 BP and Conchalí dated 700 BP) represent a distinctive lineage contributing to QH-related individuals.

We further explored the relationship between QH individuals and other present-day individuals from South America using qpGraph. We started by exploring different models for a set of present-day individuals and each QH individual in the R package ADMIXTOOLS2 [34]. Then, the parameters of the model (branch lengths and admixture proportions) were optimized using qpGraph in the software ADMIXTOOLS [31]. We evaluated different topologies based on the worst Z-score and the presence of zero-length internal branches, as well as incorporating information from our analyses about the relationship between these populations. For example, we were able to replicate the gene flow from an unsampled population (UPopA) into Mixe [3,5]. Within South America, we included representatives of the north coast of Peru (Eten), central-south Peru (Paran), and central-south Chile (Mapuche-Huilliche). Figure 4A-D summarizes the best-fit models for each QH individual. Consistent with our previous results (Fig.1-3), the different genetic affinities of each QH individual are clearly observed in the presented models. We could not find a good fit for the El Toro individual with a present-day population from Northwestern or the Cuyo region in Argentina due to the high amount of missing data after masking the present-day data for European admixture (see Methods for details on masking). However, following up on the outgroup-*f* 3 results, we included Mapuche-Huilliche from the central-south region in Chile as one of the present-day populations with the highest genetic affinity to the El Toro individual and observed that this individual is cladal to a lineage represented by present-day Mapuche-Huilliche. Since El Plomo individual showed a high genetic affinity with central-south Peru populations, we first evaluated an unadmixed model, mostly obtaining absolute Z-score values higher than 3 (Figure S12). The model that finally produced the best fit for El Plomo involves an admixture event between a lineage close to Paran and another one close to Mapuche-Huilliche. Complementary, we evaluated the best-fit qpGraph models of El Plomo or El Toro (Fig. 4A, B, and C respectively) with other ancient individuals as targets. In particular, we used the genomes from Conchalí 700BP and Pukara-6 700BP [2] as the groups that are geographically closer or share the highest genetic drift to the El Plomo, respectively. For Conchalí 700BP we used the best-fit graphs of El Plomo and El Toro. Both models provided a good fit (Figure S13A-D), but in the best-fit model of El Plomo the admixture percentage from the Paran-related lineage into the Conchali group was very low in contrast to 57% into El Plomo (0% with the full data and 3% with transversions only). We next ran a D-statistic of the form D(Huilliche, Conchalí; X, Yoruba) where X represents different present-day populations from central-south and coastal Peru. There were no values significantly different from zero, suggesting no additional contributions from X into Conchalí (Table S7). The ancient individuals that have the highest shared genetic drift with the El Plomo child are located further north, in north Chile, particularly those from the site Pukara-6. However, the best-fit qpGraph model of El Plomo failed for these individuals, finding instead a better fit with the unadmixed model (Figure 13E-H). Finally, a model of the Aconcagua individual suggested shared genetic ancestry with Eten, a present-day population representative of the north coast of Peru.

**Fig. 4.**
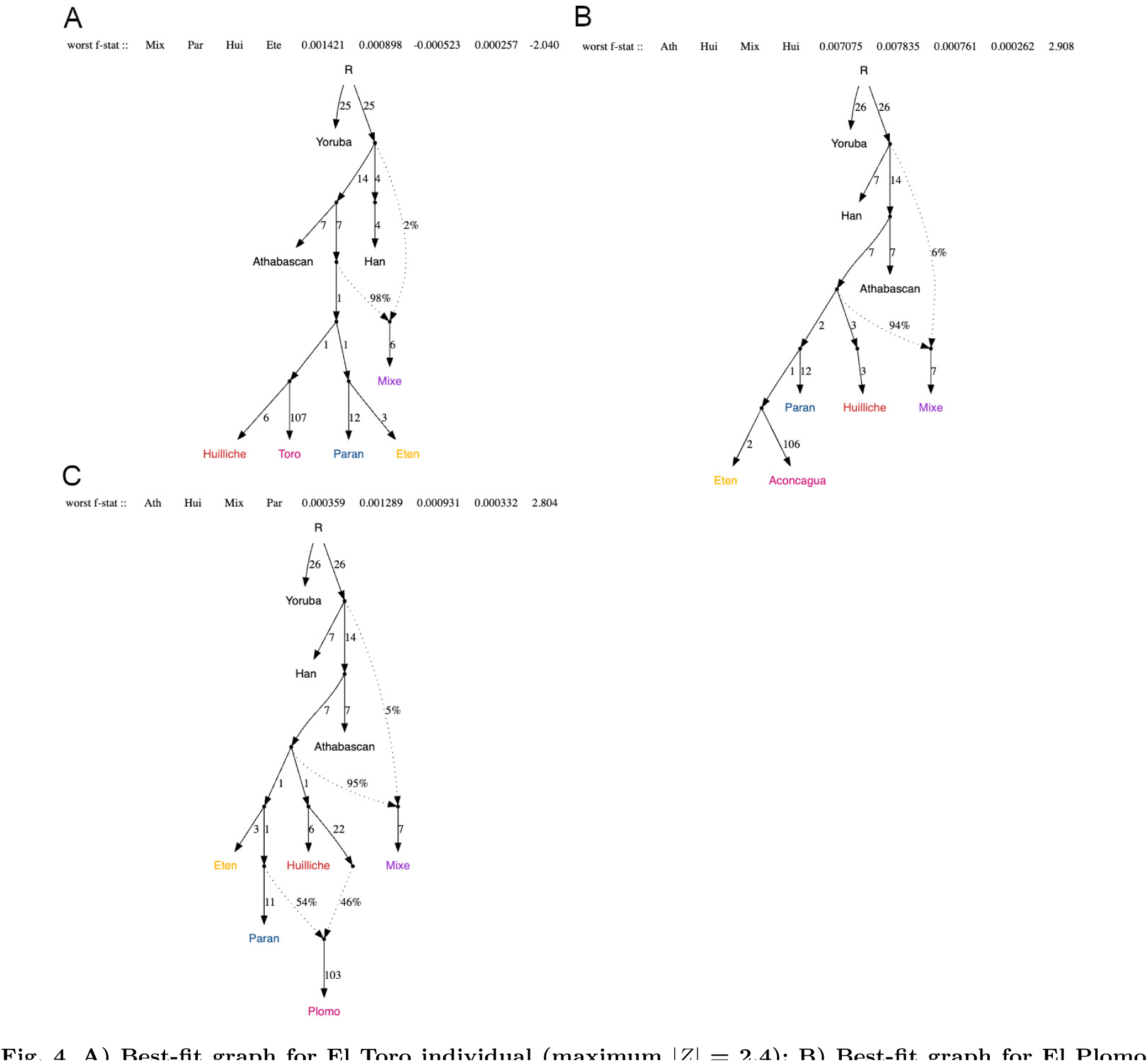
**A) Best-fit graph for El Toro individual (maximum *|Z|* = 2.4); B) Best-fit graph for El Plomo individual (maximum *|Z|* = 2.6); C) Best-fit graph for Aconcagua individual (maximum *|Z|* = 2.5).** Additional models are shown in Figure S9-S13.

We estimated the time of the admixture between the lineages represented by Paran and Mapuche-Huilliche in El Plomo using DATES [35]. We tested alternate models with modern sources representing central-south Peru to include other present-day populations than Paran that also shared high genetic drift with the El Plomo individual (Cusco, Cusco2, and Aymara). Additionally, due to potential biases introduced by recent demographic events in present-day populations, we replicated the admixture dating with ancient sources sharing the highest genetic drift with the modern sources (Paran and Mapuche-Huilliche), identified through an outgroup-*f* 3 analysis (Table S5). For Mapuche-Huilliche, we selected two alternate ancient groups, Chile Conchalí 700BP and Chile LosRieles 5100BP, while we substituted Paran with South Peru Highlands and North Peru Highlands [2]. Three pairs of sources provided a good fit by adhering to the following criteria (a) Z-score *>* 2, (b) *λ <* 200 generations, and (c) NRMSD *<* 0.7 [35]: Huilliche-Cusco, Huilliche-Aymara and Chile LosRieles 5100BP-South Peru Highlands (Table S6). The latter two pairs yielded a similar admixture time estimate of 2,000-2,600 years BCE, which suggests an admixture event between representatives of the central-south Peru and the central-south Chile lineages that predates the Inka period. On the other hand, while the Huilliche-Cusco pair yielded a younger estimate of 72 years CE, we speculate that this result may be impacted by recent drift or bottleneck in the present-day Cusco population as reported by [36].

## Discussion

The *Qhapaq hucha* ceremony has been described as a religious and political mechanism of social control over the people that the Inka conquered as they expanded into new territories. The location of these ceremonial burials at high altitudes has been linked to a sense of adoration of the Andean peaks. While this was likely a widespread sentiment across Andean societies, the Inka were probably the first to systematically explore and use the Andean peaks to their advantage as can be testified by the number of roads and structures (e.g. tambos or platforms) that were built along the Andes [17]. The individuals associated with this ceremony were mainly children or young women, buried with a rich and diverse assortment of grave goods. Textiles of different kinds, anthropomorphic and zoomorphic figurines made of different metals or *Spondylus*, feathers of different birds, and pottery are some of the most common elements found in these burials [17,37]. It is particularly interesting that the material evidence from the burial contexts was very diverse, bringing together valuable elements from the different regions of the empire or territories that were in contact with the Inka and local populations.

The diversity of the offerings and the notability of the QH ceremony itself have triggered several discussions about the origin of the buried individuals, how they were chosen, and how was their journey to their final resting place. One hypothesis is that the individuals chosen for the ceremony migrated either from the capital or from other distant areas of the empire. Alternatively, these individuals could have been part of local communities, originating from regions close to the burial sites [17,38]. It is noteworthy that the origins of these individuals, whether socio-cultural, geographical, or biological, should be considered in light of the dynamic nature of not only the Inka period but also events and population interactions across the Andes that predated the Inka period. Our genomic analyses do not favor either of the aforementioned hypotheses, reflecting a more complex scenario of human movements and interactions in the ancient Andes. This is in agreement with the varied strategies the Inka implemented across the empire to incorporate new territories and integrate new people, including the resettlement of populations and the use of intermediaries to extend its influence. The resettlement of populations was an important strategy implemented by the Inka to consolidate their position, control new regions, and organize *mitmaqkuna* laborers in relocated working camps [39]. While this was an extensive practice and it is estimated that almost a quarter or a third of the population was resettled [39,40], the number of people relocated, mode, and reason varied from region to region. Moreover, the clothing and grave goods accompanying these QH individuals are as diverse as their genetic affinities and several characteristics of the burials have contributed to evaluating their cultural affiliations and, by extension, the putative origins of the associated individuals.

The El Toro Summit is located in San Juan Province, in northwestern Argentina. Inka influence in the region has been dated to ca. 1,475 cal AD, although proof of Inka presence here is sparse compared to other areas. In fact, it has been suggested that the Inka impact in this region was associated with imperial developments on the west side of the Andes (present-day Chile) [41]. The individual buried at the summit was estimated to be 20-22 years old at the time of death. This individual is not only older than other *Qhapaq hucha* individuals, but the funerary context is, in general, less diverse, consisting of only a few textiles and no figurines or pottery [42]. While the location of the burial at high altitude (6,120 masl) constitutes one of the main features of the ceremony, the individual’s grave features and age have prompted suggestions that the El Toro individual may not have been related to the *Qhapaq hucha* ceremony or, at least, was not one of the main burials associated with the ceremony [43]. While there are only a handful of high peaks with human burials along the Andes (N = 14), they are all surrounded by several other structures of ceremonial or administrative nature, which could alternatively explain the presence of the El Toro burial [19,37]. Taken together, this implies that the El Toro individual may have either been involved in the *Qhapaq hucha* ceremony or may have served as *chaski* or messenger for the Inka, which is supported by osteological evidence of plantar keratosis that may have been caused by extensive walking, and possibly associated with a long journey or a high mobility during this individual’s life [43]. The cultural elements found in association with the individual have been linked to either the *Cuntisuyu* (western range of the Inka Empire) or *Collasuyu* (southern range of the Inka Empire). Moreover, the manufacturing and material of the clothing and other elements of the funerary contexts show a resemblance to local groups [18,43]. In concordance with the archaeological evidence, the mitochondrial haplogroup (D1j) and genome-wide diversity of the El Toro individual suggest genetic affinities with present-day and ancient populations from northwestern and Cuyo region of Argentina and central/central-south Chile, regions that were part of the southern range of the *Tawantinsuyu*.

The child from Aconcagua, located in Mendoza Province, Argentina, was a 7-8-year-old boy found at 5,250 masl. The body was wrapped in 18 pieces of textiles and accompanied by diverse grave goods, including textiles, and anthropomorphic or zoomorphic figurines made of metal and *Spondylus* [44,45]. It has been argued that the textiles and footwear of the child linked him to the Peruvian central coast, particularly the Chancay culture, evidence used in support of a coastal origin for this individual [45]. In addition, isotopic analysis of hair from the individual suggested a terrestrial diet for the year and a half before their death and, based on less conclusive results from bone collagen, a mixed diet before that [46]. Overall, there is cultural evidence suggesting affinities with groups from the Peruvian coast, but the characterization of the individual’s diet lacking a clear marine input is not concordant with this hypothesis. The genomic analyses suggest a genetic affinity of this individual with several present-day and ancient individuals from the Peruvian coast, which supports the cultural links described before and can be interpreted as evidence of long-distance movement of this individual from coastal Peru to their final burial location in Aconcagua in Argentina. These results are in agreement with a previous study [2]. However, isotopic and morphometric evidence from the Uspallata Andean Valley (Mendoza Province in Argentina, near the Aconcagua mountain) support the arrival of migrants before the Inka expansion to the area, from about CE 1,280 until 1,420 [47]. While the origin of these migrants could not be identified, this result suggests a more complicated scenario of social interactions and human migrations that pre-dates the arrival of the Inka in this region. Additional paleogenetic data from the region is needed to evaluate if the long-distance movement possibly associated with the Aconcagua child is a consequence of an earlier, pre-Inka movement or if it is part of the resettlement strategy of the Inka.

The child from El Plomo, from Santiago in Chile, is the southernmost burial associated with the *Qhapaq hucha* ceremony. This child is an 8-9-year-old boy found at an elevation of 5,400 masl [44] and dated to 1,460 CE [48]. The grave goods accompanying the child are very diverse, including bags with feathers, coca leaves, hair, nails, and deciduous teeth, in addition to three zoomorphic and anthropomorphic figurines made of metal or *Spondylus*. The clothing and ornaments (silver bracelet and headdress) suggest an association with the *Collasuyu* [44,49]. In addition, several multidisciplinary studies have been conducted to evaluate this individual’s health and cause of death, suggesting good health and evidence of trauma associated with the child’s death [48]. The genomic analyses conducted in this study suggest genetic affinities of the El Plomo child with present-day populations from central-south Andes and ancient individuals from northern Chile and south Peru highlands. Morever, our results suggest that the El Plomo child may be part of a currently unsampled genetic lineage from South America. We modeled the lineage of the El Plomo child as deriving genetically from two distinct lineages, represented by present-day populations in central-south Peru and central-south Chile. Recent research suggests that present-day populations from central-south Chile are part of a lineage splitting from other South American lineages during the Holocene [50,51]. In particular, the split between central-south Peru and central-south Chile lineages has been dated between 8,200 to 9,250 BP (7,300-6,250 BCE) [53]. This suggests that there was post-split admixture between these two lineages with the El Plomo child representing one such admixted lineage with the admixture date estimated to 2,000-2,600 years BCE (4,550-3,950 BP). When evaluating the best-fit qpGraph model of the El Plomo child against a geographically closer ancient group (Conchalí 700BP) or an individual with the highest shared genetic drift (Pukara-6 700BP), we were not able to gain further resolution into this unsampled lineage.

During the Late Period (1,400-1,536 CE) in the area near El Plomo (Mapocho basin), archaeological evidence shows the expansion of socio-cultural networks and ideological interactions, suggesting a diversification of cultural groups or units, potentiated by the expansion of the *Tawantisuyu*. Circulation of objects on a large territorial scale, new ceramic shapes and iconographies, and the emergence of stone architecture are expressions of this phenomenon. It has been proposed that the Inka deployed ideological and political incorporation strategies toward local communities, primarily through distinctive ritual congregation activities tailored to the specific characteristics of each local group [13]. However, the material archaeological evidence by itself does not allow us to distinguish between displaced non-local individuals (*mitmaqkuna*) and representatives of the *Tawantinsuyu* in the Mapocho and nearby valleys who may have been both local or non-local in origin [13]. Additional genomic data from the area is needed to evaluate the impact of these interactions on the local genetic diversity, as well as to learn more about the lineage represented by the El Plomo child and its stronger genetic affinity with geographically distant groups.

While attempting to ascertain geographical origins using paleogenomics in South America, a few critical challenges emerge. Several sub-regions and time periods are poorly or not at all represented in the genetic record, limiting a more comprehensive comparative analysis with present-day and ancient populations. This limitation may lead to simplified models of genetic similarities and origins that are clearly challenged by the human population dynamism of the Inka period and the Andes region as well as known human movements in earlier periods. There is evidence of population movements and cultural interactions long before the Inka empire, since the Middle Horizon (ca. 500–2000 BCE) or even earlier [1,47,52–54]. While previous research has suggested the establishment of genetic structure and genetic continuity more broadly in the Andes ca. 2,000 years ago [2], there is evidence of admixture and mobility throughout the region (e.g [7]). There are also indications of spatio-temporal genetic heterogeneity and movements in the genomes analyzed in this paper, some of which predate the Inka period. Moreover, while our analyses provide evidence of genetic affinities between the QH individuals and particular present-day or ancient individuals across the Andes, the presented genetic results are unable to shed light on the cultural identities or ethnicities of these individuals. Regardless of what the genetic results suggest, the final resting place of these individuals ultimately ties them to the particular territories where they were found. There was a clear intention to bury them there, with obvious implications for local communities and their histories and any speculations on these matters are beyond the realm of genetic investigations.

Finally, several concerns have been raised regarding the legal and ethical aspects of paleogenomics research in the Americas (e.g. [55–59]). The destructive nature of these studies and fragmentary archaeological records stress the importance of weighing the type and number of samples, defining clearly the particular research questions or hypotheses, and the well-planned application of current technologies. Similarly, there has been little to no record of the total number of samples taken and processed by various laboratories working on paleogenomic projects in this region versus what is eventually included in the final publications (e.g. samples that have failed to yield DNA). In this work, we aimed to evaluate the genetic diversity of individuals associated with the Inka ceremony known as Qhapaq hucha. In order to perform this research, we design the project by avoiding new destructive sampling. Instead, samples collected previously in 2005 by one of the co-authors (MM), with the aim of evaluating their genetic diversity using uniparental markers (mtDNA and Y-chromosome), were re-analyzed using next-generation sequencing technologies. Museums that previously authorized the research were re-contacted to inform about the new approach. Furthermore, we engaged in outreach activities to disseminate the results of this study and their implications prior to publication. In the process of disseminating the results of this research, we also learned more generally about some of the legal and ethical claims of Indigenous Peoples in Argentina concerning their Ancestors, and mostly associated with fights for their recognition and ancestral rights [60–63]. We stress the importance of dialogues with local researchers and Indigenous communities not only to seek approval for research but also to learn about the history of the region and to weigh the consequences of our research and narratives to present-day communities.

Overall, on a local scale, this study contributes novel results to our growing understanding of the nature of the *Qhapaq hucha* ceremony, with a focus on the genetic origins of the buried individuals. More broadly, it expands our knowledge of human genetic variation in South America prior to European colonization with the identification of a previously unsampled lineage as well as pondering the evidence or expectations of gene flow in the focal time period. Future archaeogenomics research implementing appropriate ethical and community-engaged strategies will provide greater resolution on the bio-cultural dynamics in the region.

## Methods

### Data generation

No new samples were collected in this study. Whole genome data was generated from samples collected in 2005, initially studied using only mitochondrial DNA markers (D-loop). Museums were notified of the new study aims and methods and they agreed with implementing newer methodologies. DNA extractions were performed from tissue (muscle) or bone samples, using the protocols described in [64–66]. Double-stranded DNA libraries were built following the standard protocol from Meyer and Kitcher, 2010 and sequenced on an Illumina MiSeq and NextSeq. All laboratory work and sequencing were performed at the Faculty of Medicine of Universidad de Chile. Pre-PCR work was carried out in a facility dedicated to the analysis of aDNA samples, which is isolated from laboratories working with DNA from present-day samples. This facility has positive air pressure (HEPA-filtered and UV-treated airflow) and UV lamps on all working surfaces. Samples and reagents were manipulated under a laminar flow cabinet and using disposable sterile plastics and consumables. Raw sequencing reads were mapped to the human genome reference build hg19 and the mitochondrial reference rCRS using BWA (aln and seed disable). Unmapped reads and those reads with mapping quality below 30 were removed using samtools. PCR duplicates were identified and removed using Picard MarkDuplicates. Finally, we used GATK for indel realignment and samtools calmd to generate MD tags. Pseudo-haploid calls were generated using ATLAS for all four ancient individuals (Aconcagua from BAM) and for all ancient genomes with available shotgun sequencing data (see Table S3). For those ancient genomes with only published SNP data (1240k panel), files were downloaded in eigenstrat format (see Table S3).

### Reference dataset

We compiled a database of almost 3,400 worldwide present-day individuals (Table S2) and 296 ancient individuals (Table S3) from South America. Since most present-day data was generated using different SNP arrays (Axiom LAT1 and Human Origin Affymetrix), this data was phased and imputed by array using TOPMed Michigan Imputation Server. ADMIXTURE analysis was performed before and after imputation to evaluate any changes in their global ancestry (Figure S15). We kept the sum of positions from both arrays and the 1240k positions from the aDNA enrichment panel, for a total of 1.6M positions. Present-day South American individuals with more than 99% Native American ancestry (according to K=3 in ADMIXTURE) were used as references together with individuals with more than 99% European or African genetic ancestry in order to estimate local ancestry using RFMix v2 with the parameters described in [67]. Non-Native American genetic ancestry was masked and, unless otherwise indicated, all analyses were performed using this masked dataset.

### Analyses

Principal Component Analysis was performed using smartpca from Eigensoft [30], with ancient individuals projected using lsqproj = YES. All f-statistic-based analyses (outgroup-*ƒ* 3, D-statistic, qpWave, and qpGraph) were performed using the software ADMIXTOOLS [31]. For the qpGraph analysis, we started by exploring different models for a set of present-day individuals and each QH individual in the R package Admixtools2 [34]. Then, the parameters of the model (branch lengths and admixture proportions) were optimized using qpGraph in the software ADMIXTOOLS [31]. The set of present-day populations includes Yoruba and Han from 1000G dataset, Athabascan [66], Mixe [68], Eten, Paran [36], and Mapuche-Huilliche [69]. The qpWave analysis was implemented in the R package Admixtools2 [34]. Results were plotted using Rstudio (http://www.rstudio.com/) and custom scripts.

### Availability of data and materials

Data will be available for download through the European Nucleotide Archive (ENA) (accession no. xxxxx).

## Authors’ contributions

CDC, MC and MM designed the research; RV and MM provided funds and resources to perform the research; CDC, DC and MM performed the research and analyzed the data; CDC, CC, MR, MC and MM contributed to interpreting results; CDC, CC, MR and MM wrote the paper with input from all co-authors.

## Acknowledgements.

We would like to thank several institutions and individuals contributing to this research: Instituto de Investigaciones Arqueológicas y Museo ”Prof. Mariano Gambier”; Catalina Teresa Michieli; Museo Nacional de Historia Natural, Santiago, Chile; and Illumina Inc. We are grateful for the support provided by Mariela Rodriguez and Marina Sardi regarding legal and ethical matters in Argentina, and to Nadia Gomez, authority of the Warpe Community in the Cuyum Territory and Dra. Carina Jofre, CONICET researcher and member of the same Warpe Community, for sharing their knowledge regarding the research of ancestral human remains and their critical view of our work. Finally, we thank Lumila Menendez for her valuable comments to the manuscript.

## Supplementary information

This article has supplementary figures and tables.

