## Supplementary figures and tables for "The genomic and cultural diversity of the Inka Qhapaq hucha ceremony in Chile and Argentina"

**This PDF file includes:**

Figures S1 to S15
Tables S1 to S7
SI References

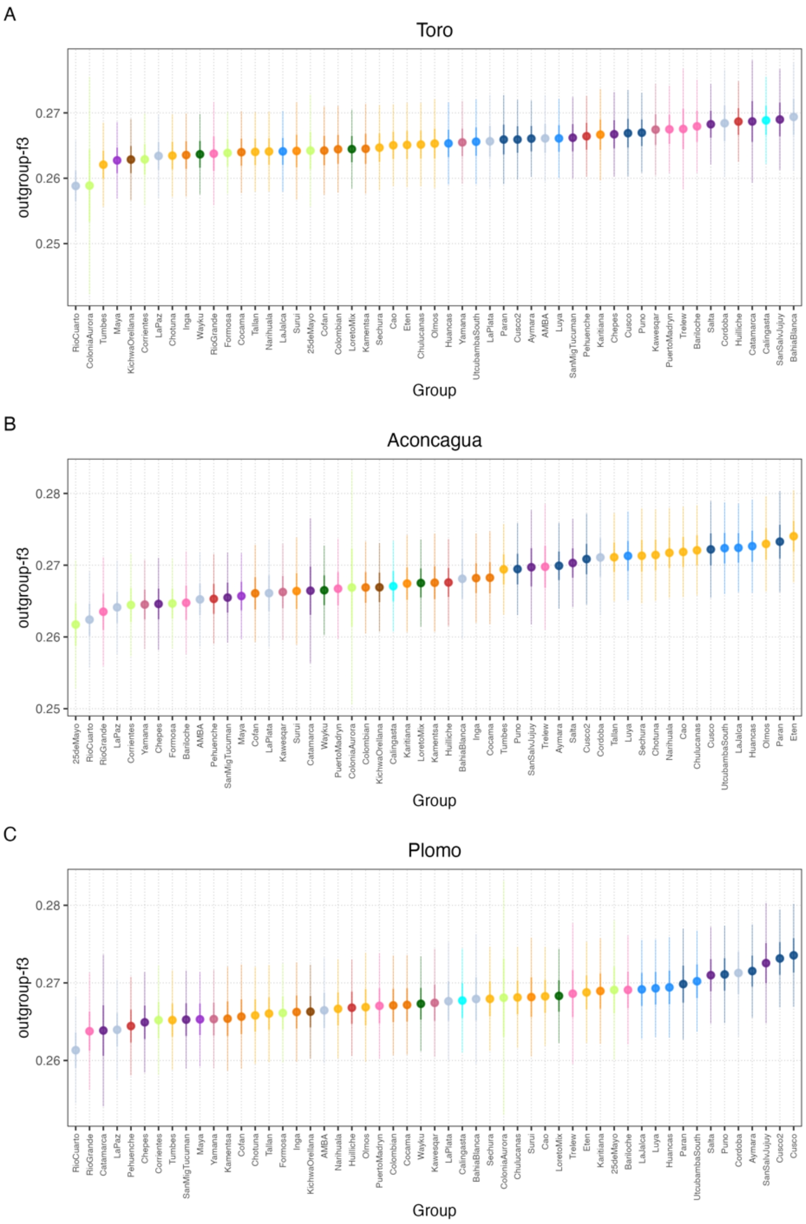

**Fig. S1. Extended outgroup-ƒ3 analysis.** The analysis was performed with the software admixtools (1) of the form ƒ3(QH, X; Outgroup), where QH represents each of the three ancient genomes associated with the Qhapaq hucha ceremony and X represents different present-day populations from South America.

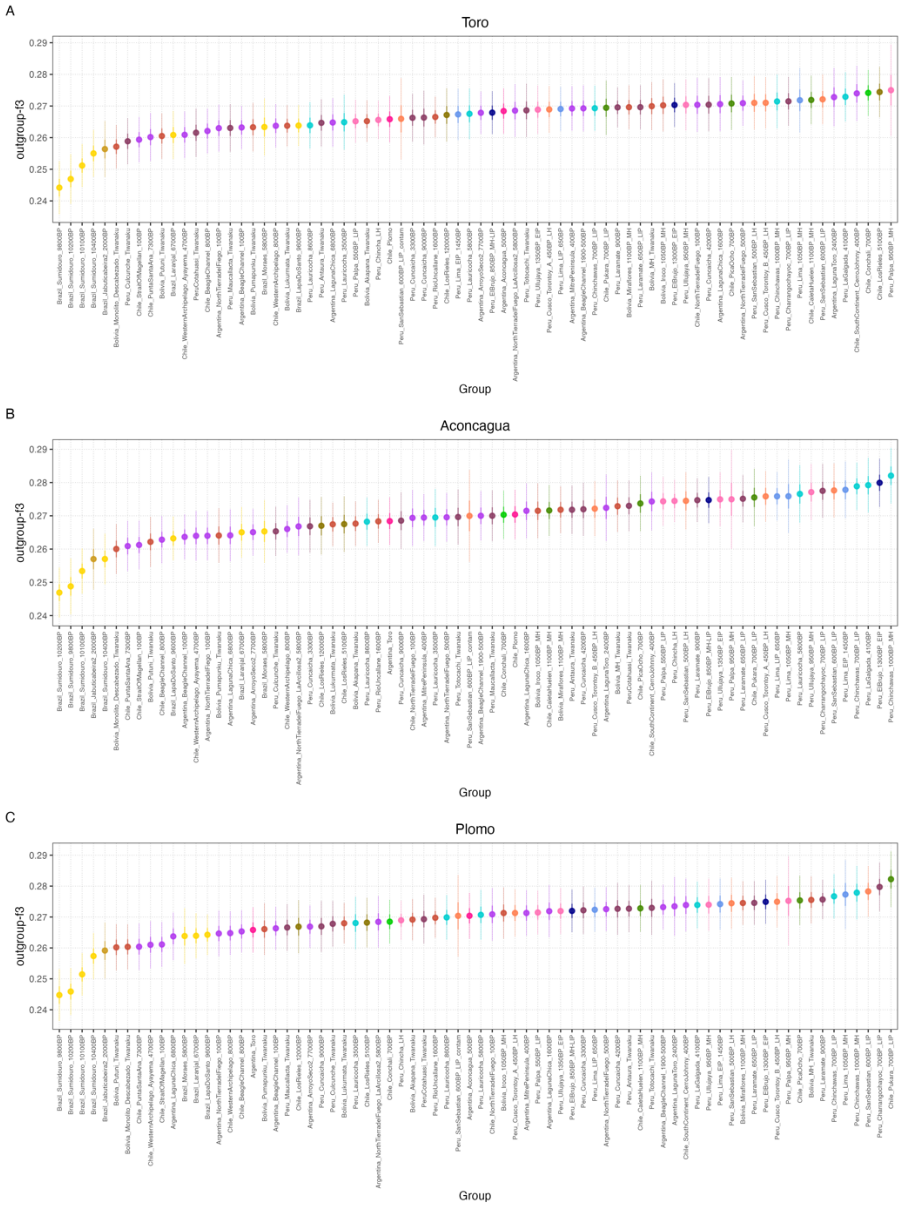

**Fig. S2. Outgroup-ƒ3 analysis.** The analysis was performed with the software admixtools (1) of the form ƒ3(QH, X; Outgroup), where QH represents each of the three ancient genomes associated with the Qhapaq hucha ceremony and X represents different ancient groups along South America.

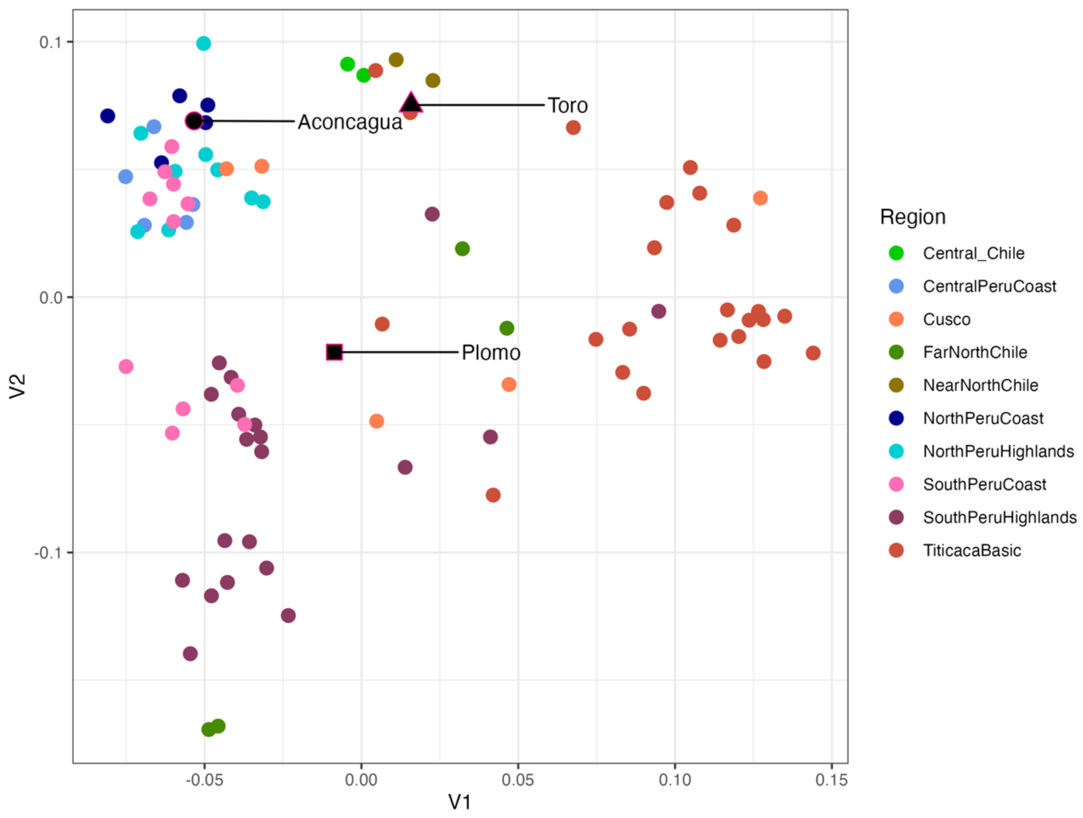

**Fig. S3. Multidimensional scaling (MDS) based on outgroup-ƒ3.** An outgroup-ƒ3 analysis was performed between pairs of ancient individuals from South America using qp3pop in Admixtools (1). Only pairs with more than 5,000 overlapping positions and values converted to distance (1-outgroup ƒ3). An MDS was calculated using *cmdscale* in R.

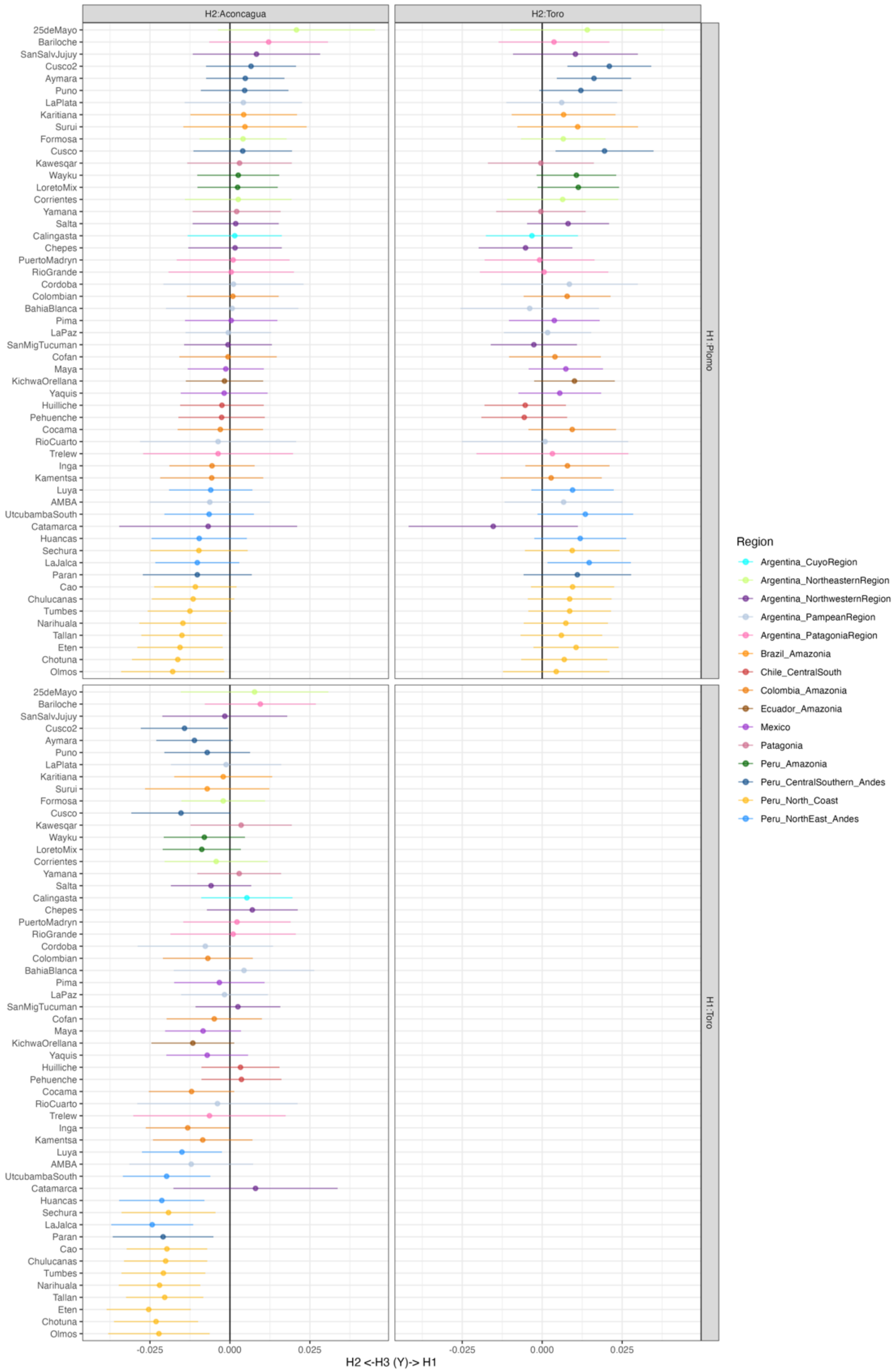

**Fig. S4. Extended D-statistic of the form D(Qh1, Qh2; X; Outgroup).** QH1 and QH2 represent pairs of Qhapaq hucha individuals and X represents different present-day populations from South America. The error bars represent 3 SDs. The analysis was performed using the software admixtools (1).

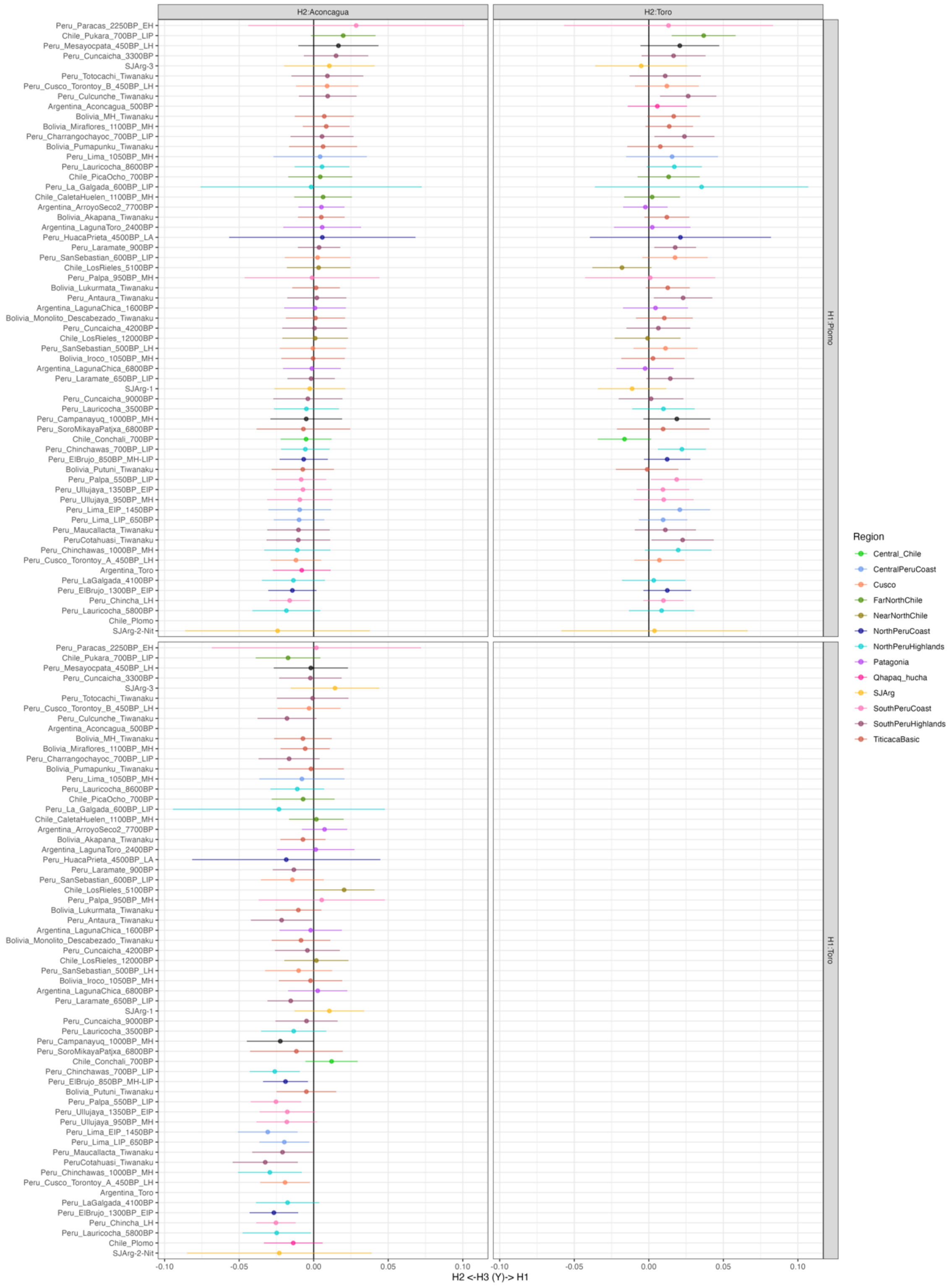

**Fig. S5. D-statistic of the form D(Qh1, Qh2; X; Outgroup)**. QH1 and QH2 represent pairs of Qhapaq hucha individuals and X represents different ancient groups from South America. The error bars represent 3 SDs. The analysis was performed using the software admixtools (1).

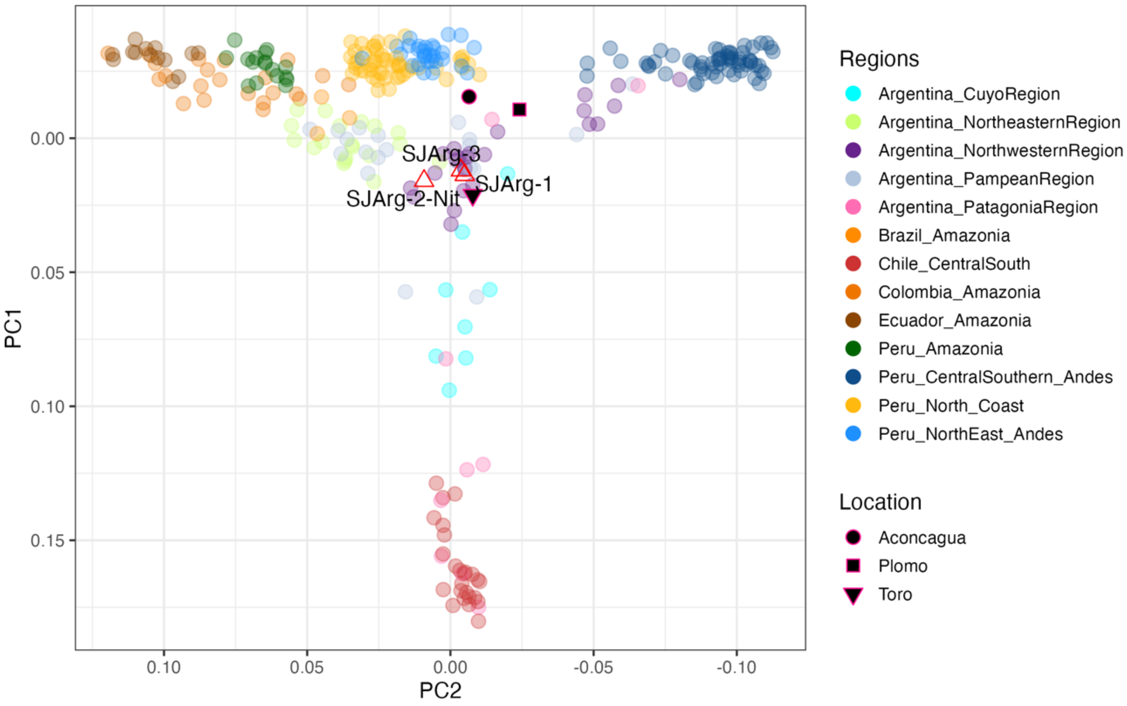

**Fig. S6. PCA including ancient individuals from San Juan, projected onto PCs 1 and 2 estimated with present-day populations that are color-coded by region** (as in Fig. 1 of the main text).

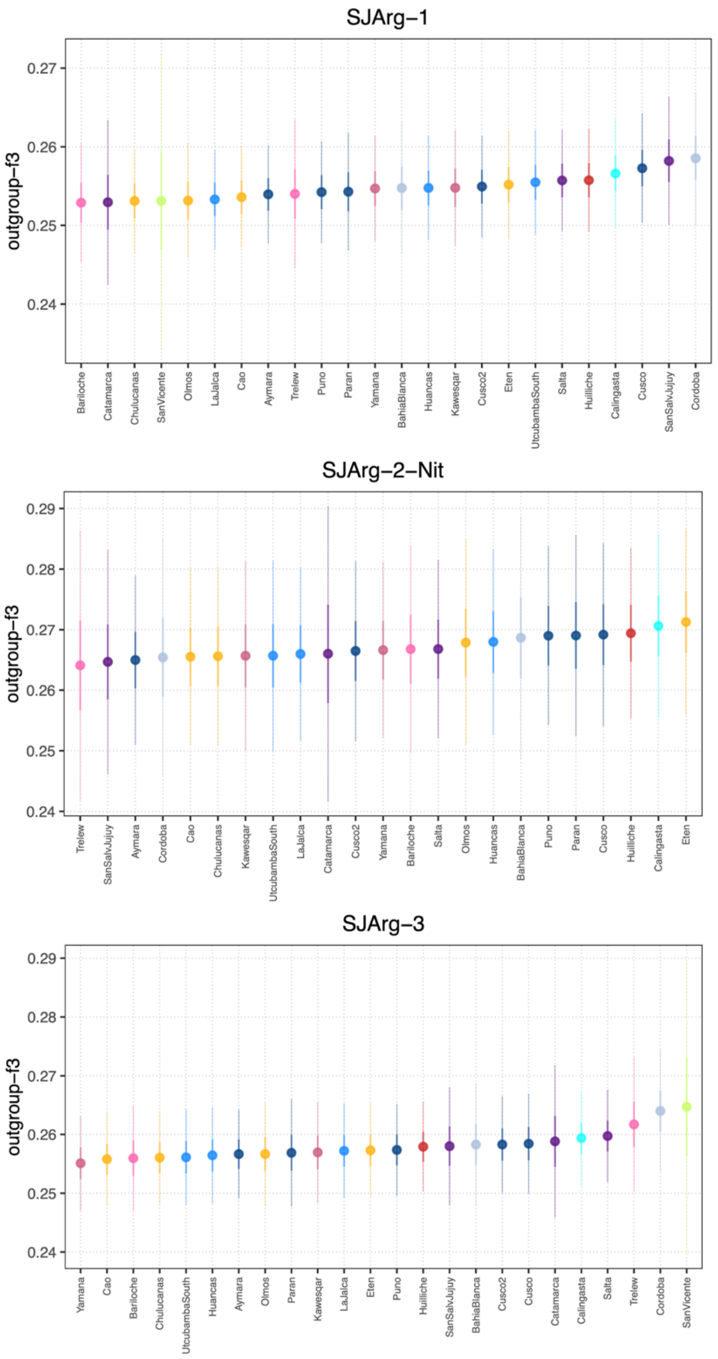

**Fig. S7. Outgroup-ƒ3 analysis.** The analysis was performed with the software admixtools (1) of the form ƒ3(SJ, X; Outgroup), where SJ represents each of the three ancient genomes from San Juan and X represents different present-day populations along South America.

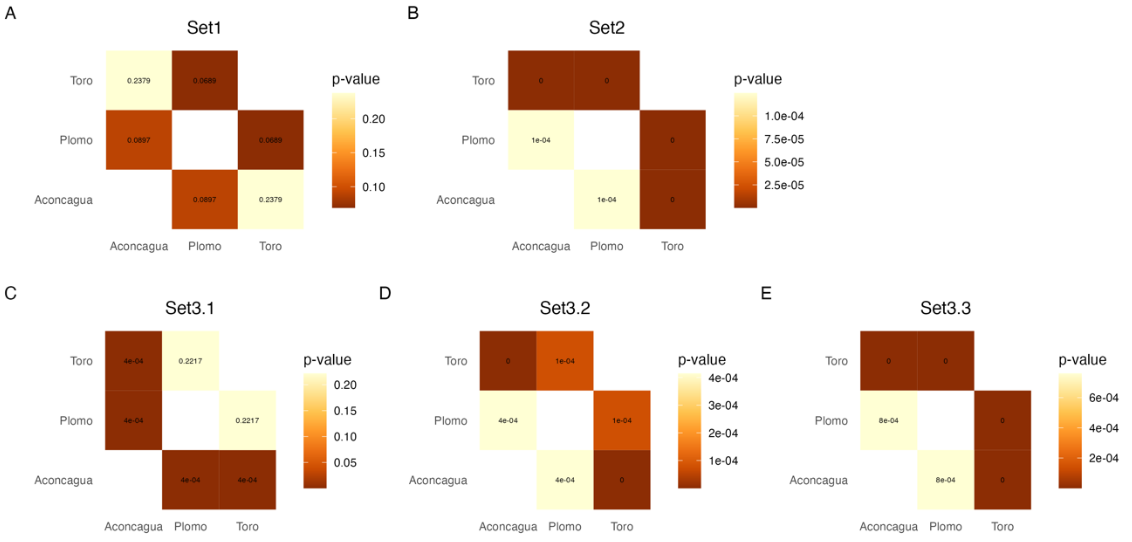

**Fig. S8. Pairwise p-values between pairs of QH individuals and different sets of "right populations".** A) set1; B) set2; C) set3_1; D) set3_2; E) set3_3. Each set has a different list of "right populations", as shown in Table S3. This test was implemented to evaluate if QH can be modelled similarly using a set of right populations. We use a threshold for p-values of 0.05 (p-value < 0.05, qpWave does not support cladality).

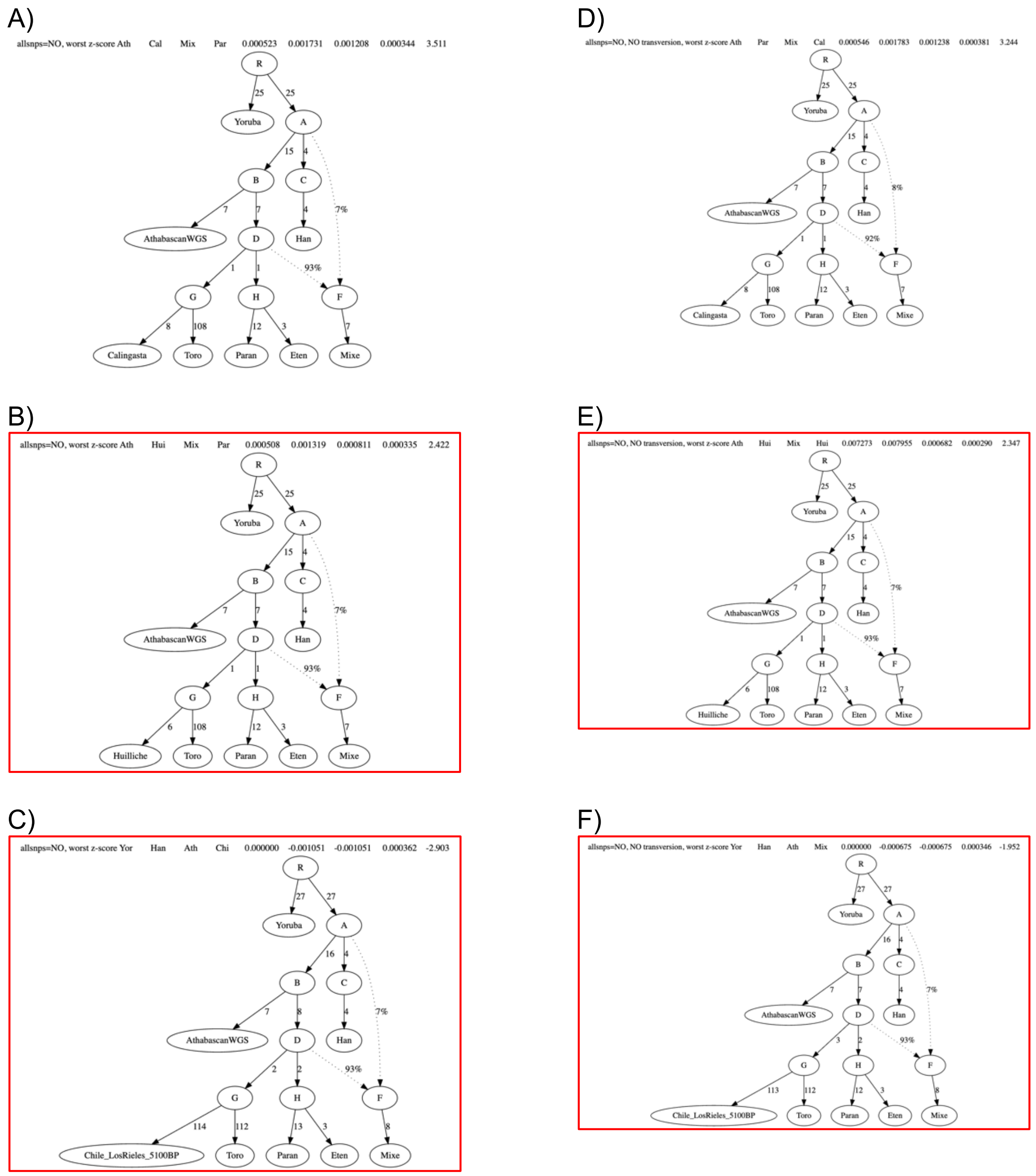

**Fig. S9. Alternative qpGraph models for El Toro.** Highlighted in red models with a good fit.

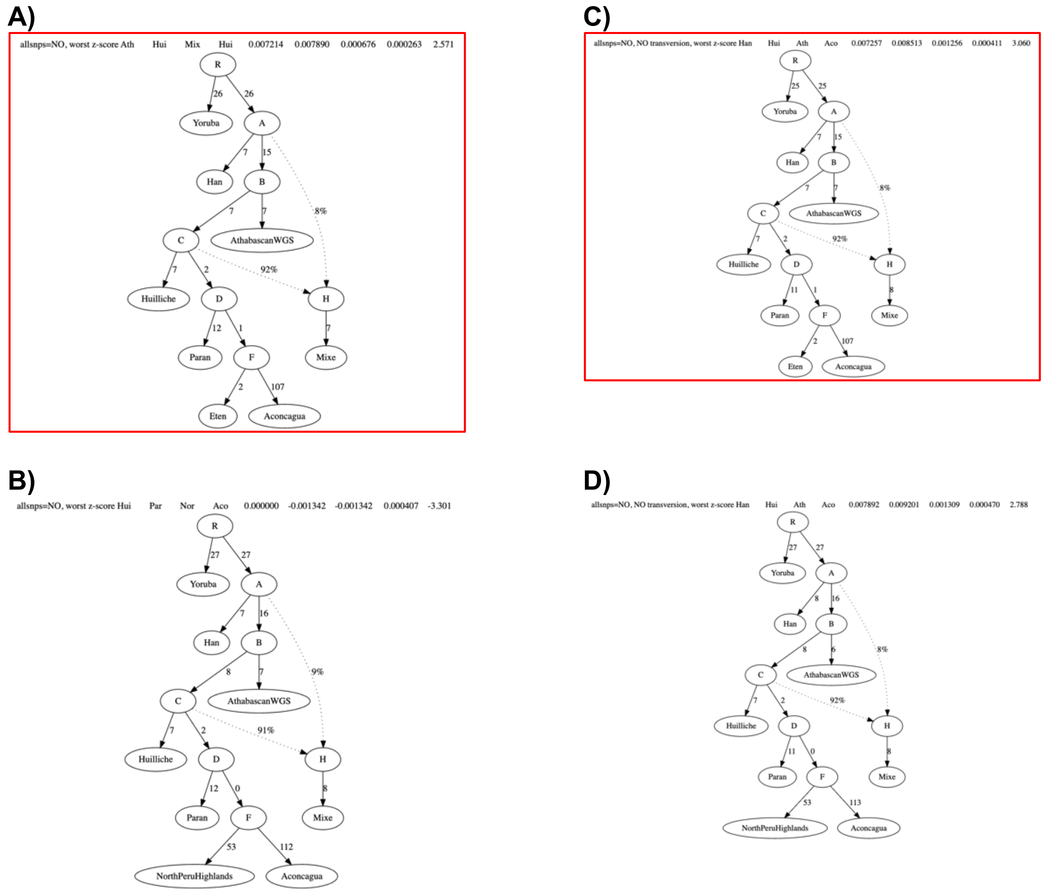

**Fig. S10. Alternative qpGraph models for Aconcagua.** Highlighted in red models with a good fit.

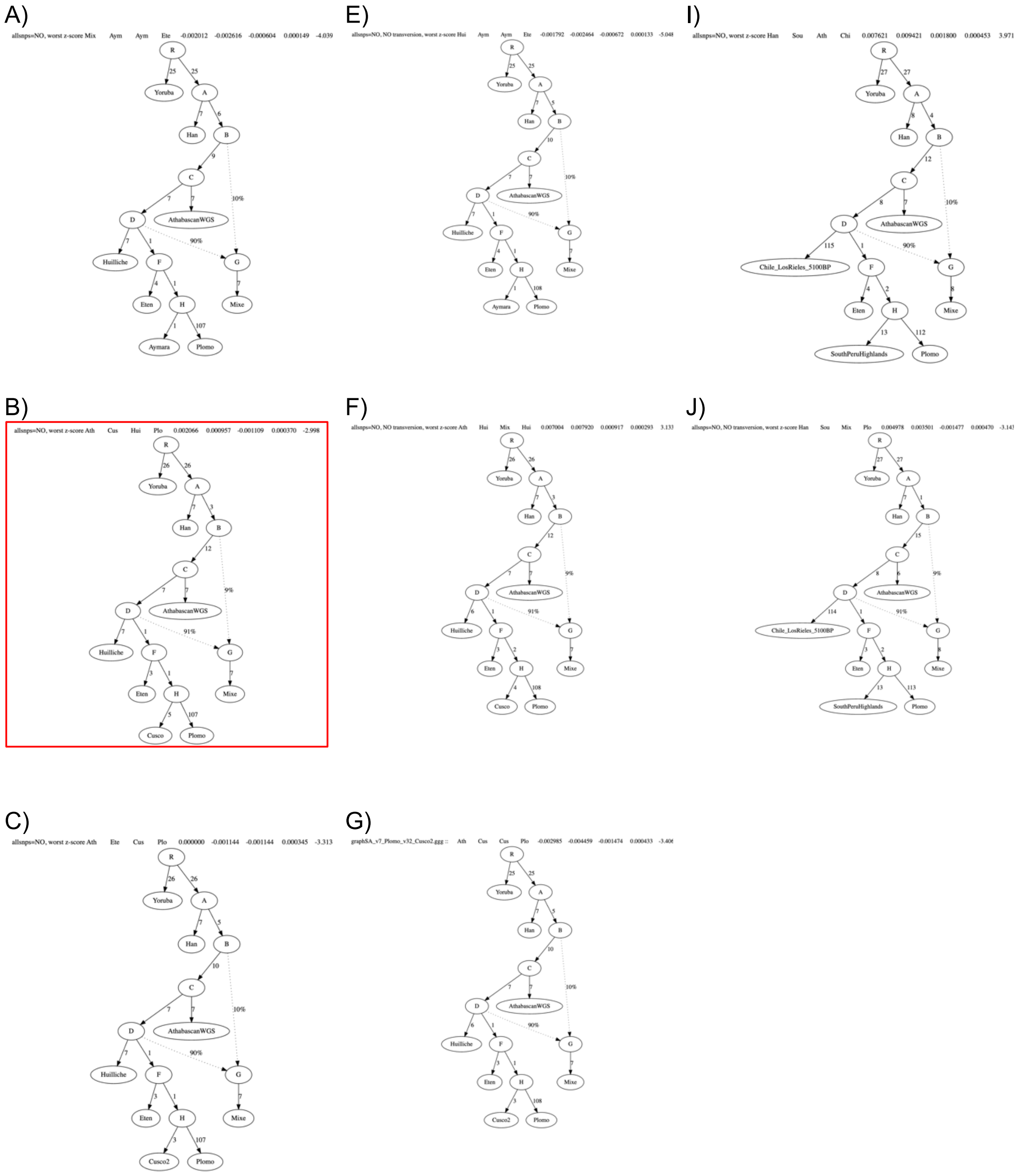
A)
allsnps=NO, worst z-score Mix Aym Aym Ete -0.002012 -0.002616 -0.000604 0.000149 -4.039
E)
allsnps=NO, NO transversion, worst z-score Hui Aym Aym Ete -0.001792 -0.002464 -0.000672 0.000133 -5.048
I)
allsnps=NO, worst z-score Han Sou Ath Chi 0.007621 0.009421 0.001800 0.000453 3.971
B)
allsnps=NO, worst z-score Ath Cus Hui Plo 0.002066 0.000957 -0.001109 0.000370 -2.998
F)
allsnps=NO, NO transversion, worst z-score Ath Hui Mix Hui 0.007004 0.007920 0.000917 0.000293 3.133
J)
allsnps=NO, NO transversion, worst z-score Han Sou Mix Plo 0.004978 0.003501 -0.001477 0.000470 -3.143
C)
allsnps=NO, worst z-score Ath Ete Cus Plo 0.000000 -0.001144 -0.001144 0.000345 -3.313
G)
graphSA_v7_Plomo_v32_Cusco2.ggg :: Ath Cus Cus Plo -0.002985 -0.004459 -0.001474 0.000433 -3.406
R
Yoruba
Han
AthabascanWGS
Huilliche
Eten
Aymara
Cusco
Cusco2
Plomo
Mixe
Chile_LosRieles_5100BP
SouthPeruHighlands

D)

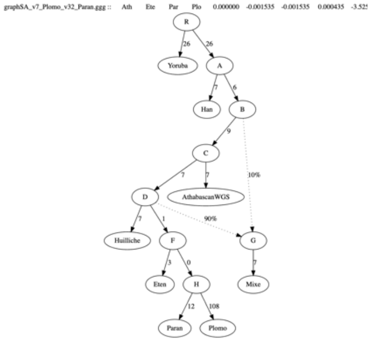

H)

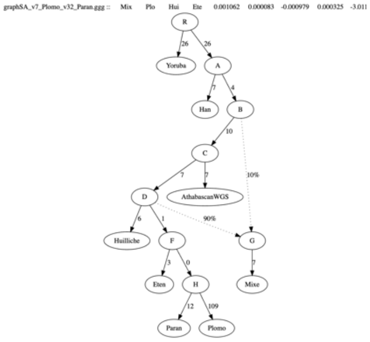

**Fig. S11. Alternative qpGraph models for El Plomo (unadmixed).** Highlighted in red models with a good fit.

A)

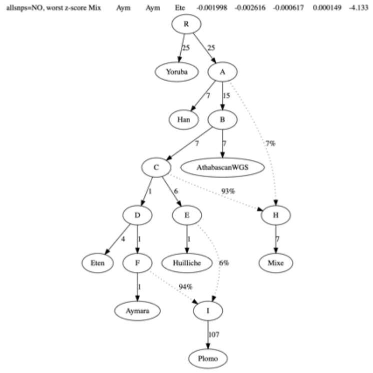

E)

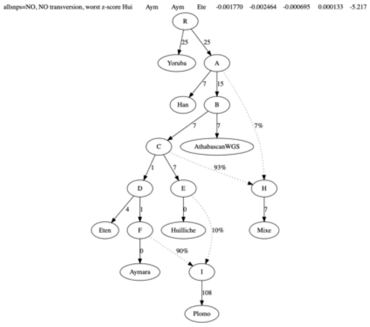

I)

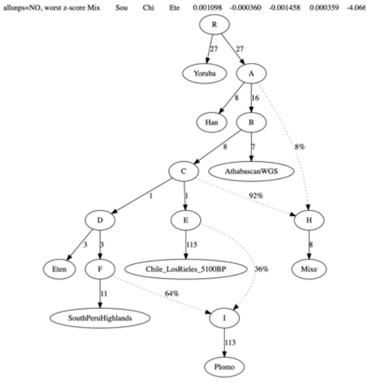

B)

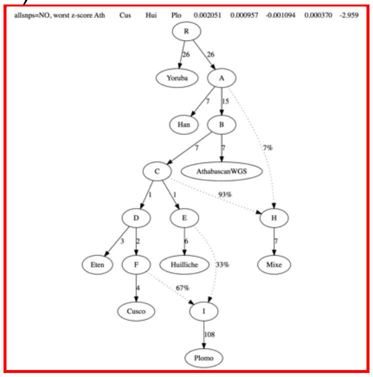

F)

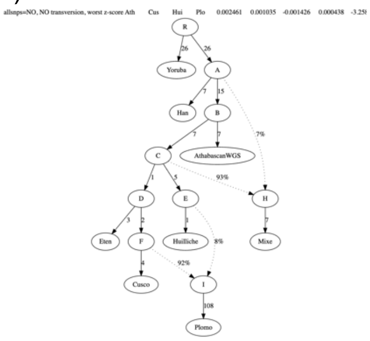

J)

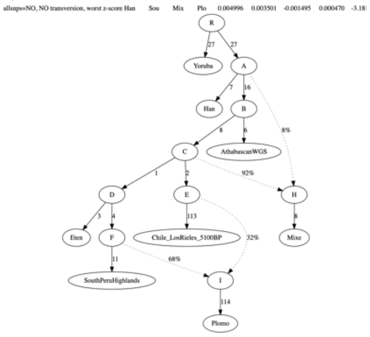

C)

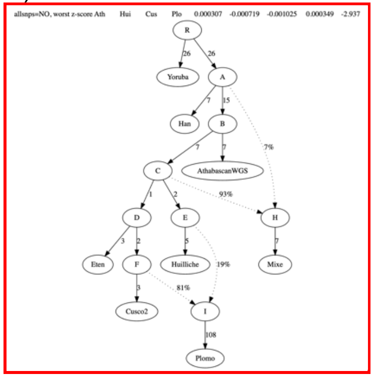

G)

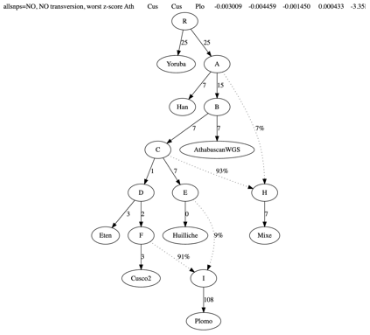

K)

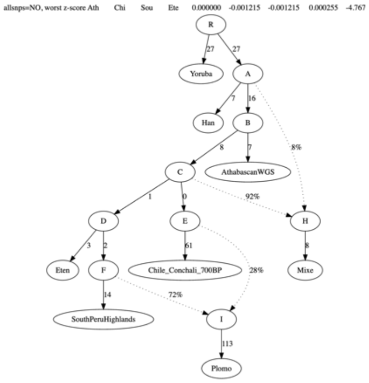

D)

H)

L)

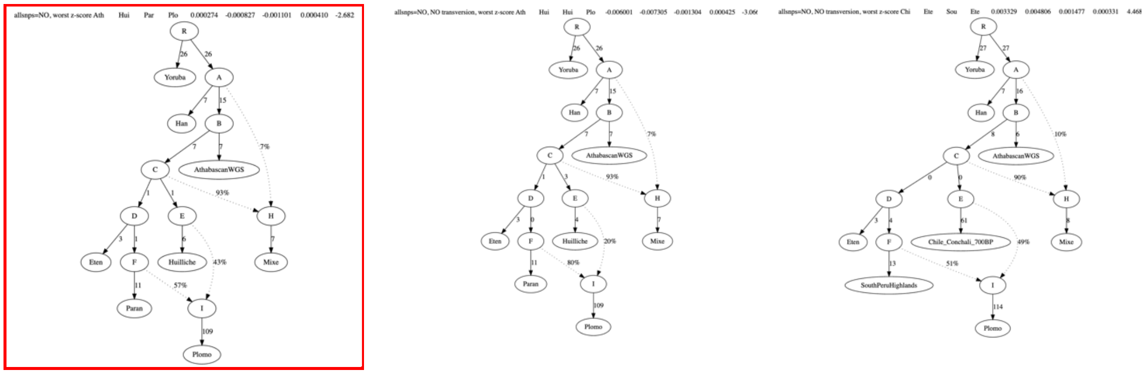

**Fig. S12. Alternative qpGraph models for El Plomo (admixed).** Highlighted in red models with a good fit.

A)

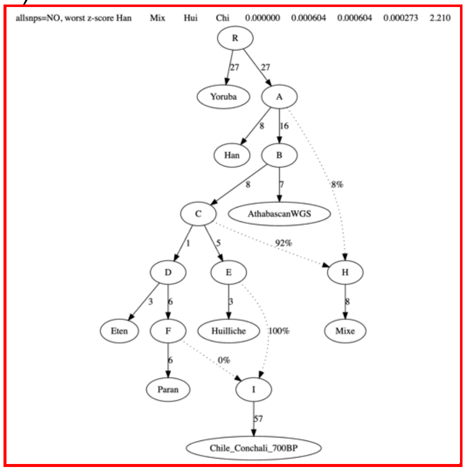
allsnps=NO, worst z-score Han Mix Hui Chi 0.000000 0.000604 0.000604 0.000273 2.210
Yoruba
Han
AthabascanWGS
Eten
Paran
Huilliche
Mixe
Chile_Conchali_700BP

B)

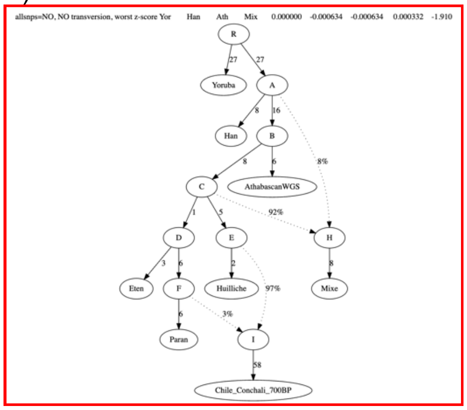
allsnps=NO, NO transversion, worst z-score Yor Han Ath Mix 0.000000 -0.000634 -0.000634 0.000332 -1.910
Yoruba
Han
AthabascanWGS
Eten
Paran
Huilliche
Mixe
Chile_Conchali_700BP

C)

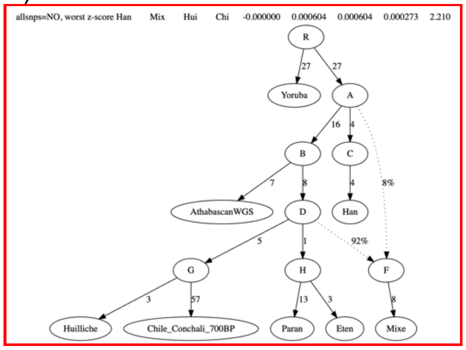
allsnps=NO, worst z-score Han Mix Hui Chi -0.000000 0.000604 0.000604 0.000273 2.210
Yoruba
AthabascanWGS
Han
Huilliche
Chile_Conchali_700BP
Paran
Eten
Mixe

D)

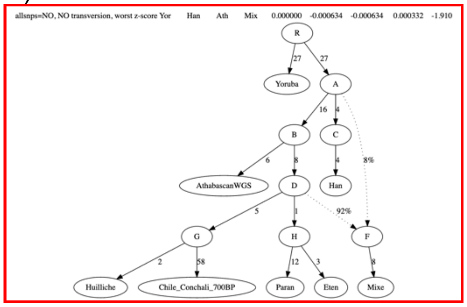
allsnps=NO, NO transversion, worst z-score Yor Han Ath Mix 0.000000 -0.000634 -0.000634 0.000332 -1.910
Yoruba
AthabascanWGS
Han
Huilliche
Chile_Conchali_700BP
Paran
Eten
Mixe

E)

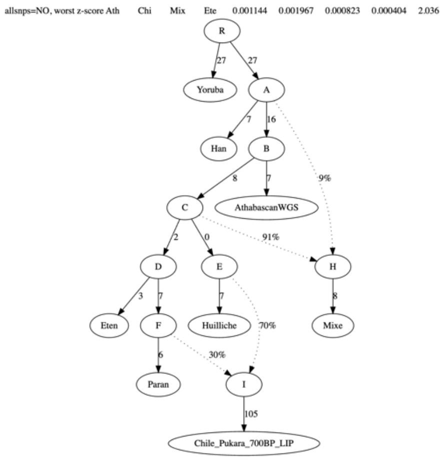
allsnps=NO, worst z-score Ath Chi Mix Ete 0.001144 0.001967 0.000823 0.000404 2.036
Yoruba
Han
AthabascanWGS
Eten
Paran
Huilliche
Mixe
Chile_Pukara_700BP_LIP

F)

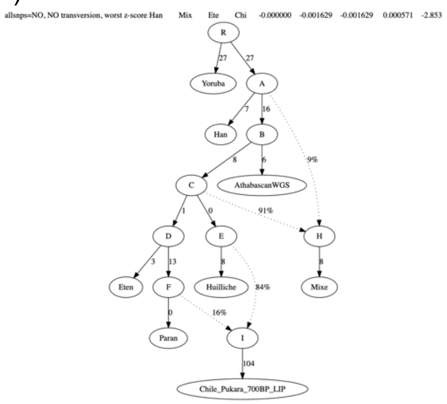
allsnps=NO, NO transversion, worst z-score Han Mix Ete Chi -0.000000 -0.001629 -0.001629 0.000571 -2.853
Yoruba
Han
AthabascanWGS
Eten
Paran
Huilliche
Mixe
Chile_Pukara_700BP_LIP

G)

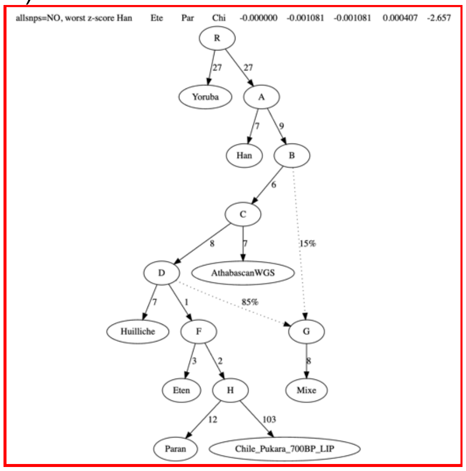

H)

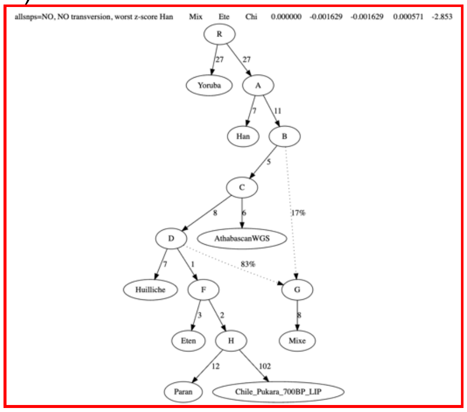

**Figure S13: Alternative qpGraph models for Conchalí_700BP (A-D) and Pukara-6_700BP (E-H)**. Highlighted in red models with a good fit.

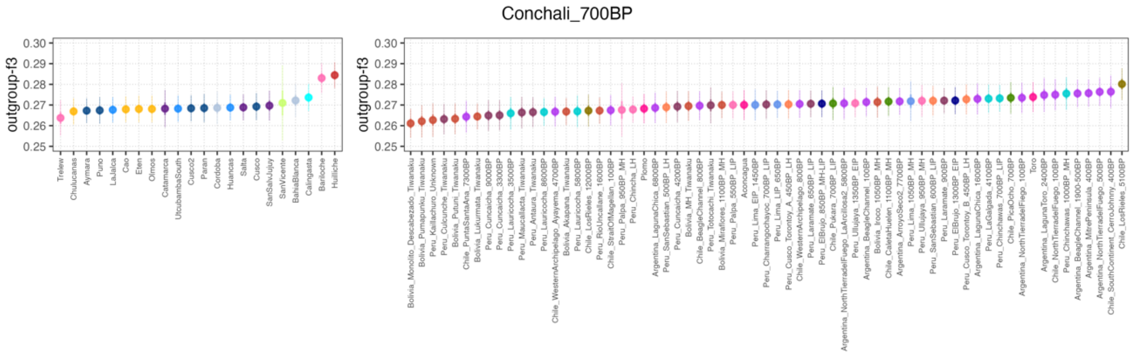

**Figure S14: Outgroup-$f3$ analysis**. The analysis was performed with the software admixtools (Patterson et al., 2012) of the form $f3$(Conchali, X; Outgroup), where X represents different present-day and ancient groups along South America.

A) Global ancestry before imputation in TOPMed and RFMIX analysis.

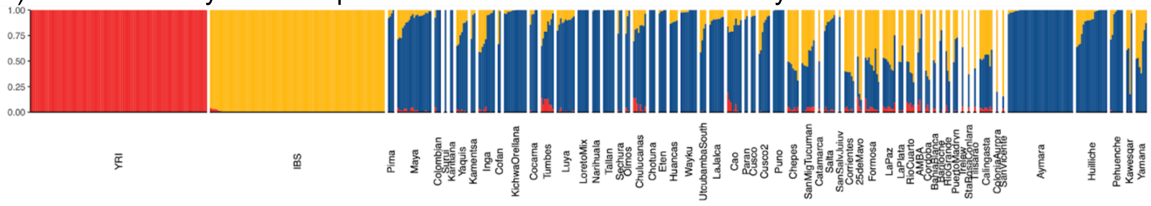

B) Global ancestry after imputation in TOPMed and RFMIX analysis.

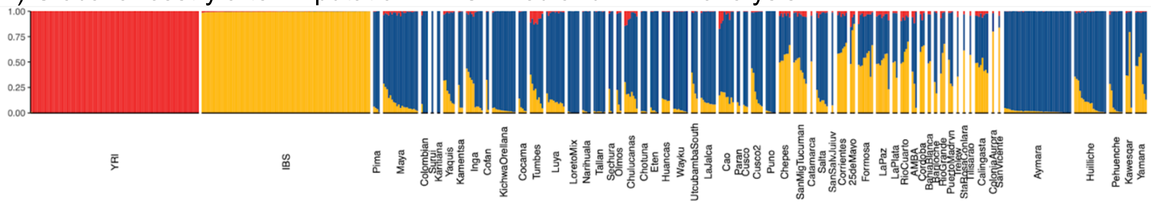

C) Correlation between global ancestry values of individuals before and after imputation and RFMIX. AMR=American (continental) ancestry-like; EUR=European ancestry-like; AFR=African ancestry-like. adm=global ancestry before imputation using TOPMed and RFMIX. rfmix=global ancestry after imputation using TOPMed and RFMIX. This value was summarised using local ancestry results from RFMIX.

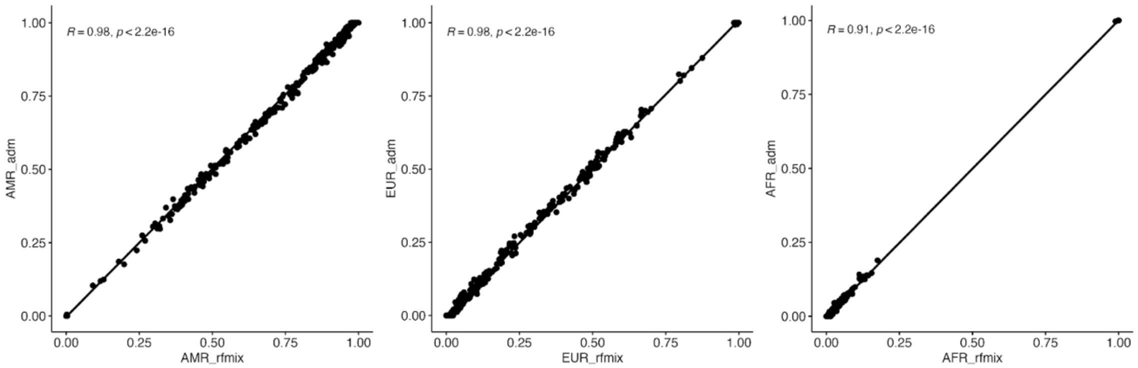

**Figure S15: ADMIXTURE analysis of reference panel before and after imputation and RFMIX analysis**.

**Table S1. Sequencing statistics.**

| Summit | DoC* | Total reads | Reads mapping | Unique reads | Endogenous percentage | Average read lenght |
|---|---|---|---|---|---|---|
| El Toro | 3.6 | 260444415 | 178548729 | 169932589 | 65.25 | 81.97 |
| El Plomo | 2.2 | 285731769 | 197857861 | 191287203 | 66.95 | 84.07 |

**Table S2. Summary of genome-wide data from present-day South American populations.**

| Study | Number of individuals | Array |
|---|---|---|
| Raghavan et al., 2015* | 2 | WGS |
| Barbieri et al., 2019 | 176 | Affymetrix Human Origins |
| de la Fuente et al., 2018 | 61 | Axiom LAT1 |
| Luisi et al., 2020 | 87 | Axiom LAT1 |
| Lindo et al., | 48 | WGS |
| HGDP** | 929 | WGS |
| 1000G*** | 1902 | WGS |
| SGDP**** | 279 | WGS |

* Only Athabascan individuals
** Bergström et al., 2020
*** Genomes Project C, et al.
**** Mallick et al, 2016

**Table S3. Summary of genome-wide data from ancient individuals from the Americas.**

| Study | Number of individuals | Data type |
|---|---|---|
| Pedersen et al., 2022 | 3 | WGS |
| Popovic et al., 2021 | 17 | 1240k capture |
| Nakatsuka et al., 2020a | 65 | 1240k capture |
| Nakatsuka et al., 2020b | 20 | 1240k capture |
| Bongers et al., 2020 | 6 | WGS |
| Posth et al., 2018 | 49 | 1240k capture |
| Moreno-Mayar et al., 2018a | 2 | WGS |
| Moreno-Mayar et al., 2018b | 15 | WGS |
| Lindo et al., 2018 | 7 | WGS |
| Scheib et al., 2018 | 91 | WGS |
| Raghavan et al., 2015 | 23 | WGS |
| Rasmussen et al., 2014 | 1 | WGS |

**Table S4. Set of "right" populations for qpWave analysis.**

| Set1* | Set2 | Set3_1 | Set3_2 | Set3_3 |
|---|---|---|---|---|
| USA_CA_Chumash | USA_Anzick | Chile_CaletaHuelen_1100BP_MH | Chile_CaletaHuelen_1100BP_MH | Chile_CaletaHuelen_1100BP_MH |
| USA_CA_SanFranciscoBay | USA_CA_Early_SanNicolas | Bolivia_MH_Tiwanaku | Bolivia_MH_Tiwanaku | Bolivia_MH_Tiwanaku |
| USA_CA_Early_SanNicolas | Mixe | Peru_ElBrujo_1300BP_EIP | Peru_ElBrujo_1300BP_EIP | Peru_ElBrujo_1300BP_EIP |
| USA_CA_LSCI | Brazil_LapaDoSanto_9600BP | Peru_Lima_LIP_650BP | Peru_Lima_LIP_650BP | Peru_Lima_LIP_650BP |
| Athabascan | Argentina_ArroyoSeco2_7700BP | Peru_Ullujaya_1350BP_EIP | Peru_Ullujaya_1350BP_EIP | Peru_Ullujaya_1350BP_EIP |
| Russia_MA1 | Bolivia_Miraflores_1100BP_MH | Peru_Chinchawas_1000BP_MH | Peru_Chinchawas_1000BP_MH | Peru_Chinchawas_1000BP_MH |
| USA_Anzick | Peru_Cuncaicha_4200BP | Peru_Laramate_900BP | Peru_Laramate_900BP | Peru_Laramate_900BP |
| Han | Peru_ElBrujo_1300BP_EIP | | Chile_LosRieles_5100BP | Chile_Conchali_700BP |
| Papuan | Peru_Chinchawas_1000BP_MH | | | |
| Zapotec | Peru_Lauricocha_5800BP | | | |
| Mixtec | Peru_Lauricocha_8600BP | | | |
| Maya | Peru_Lima_EIP_1450BP | | | |
| Mixe | Peru_Lima_LIP_650BP | | | |
| | Peru_Ullujaya_1350BP_EIP | | | |
| | Chile_CaletaHuelen_1100BP_MH | | | |
| | Chile_Conchali_700BP | | | |
| | Chile_LosRieles_5100BP | | | |

* Similar to Posth et al., 2018.

**Table S5. Outgroup-ƒ3 analysis of the form ƒ3(Huilliche/Paran, X; Yoruba)**. X represents different ancient individuals from South America.

| Pop1 | Pop2 | Outgroup | outgroup-ƒ3 | se | Z-score | snps |
|---|---|---|---|---|---|---|
| Huilliche | Chile_Conchali_700BP | Yoruba | 0.28443 | 0.002087 | 136.316 | 822947 |
| Huilliche | Chile_SouthContinent_CerroJohnny_400BP | Yoruba | 0.277629 | 0.002356 | 117.857 | 449874 |
| Huilliche | Chile_LosRieles_5100BP | Yoruba | 0.277567 | 0.002264 | 122.594 | 705239 |
| Huilliche | Argentina_BeagleChannel_1900-500BP | Yoruba | 0.276704 | 0.002118 | 130.651 | 871542 |
| Huilliche | Argentina_NorthTierradelFuego_500BP | Yoruba | 0.276599 | 0.002086 | 132.591 | 852124 |
| Huilliche | Chile_NorthTierradelFuego_100BP | Yoruba | 0.276293 | 0.002271 | 121.644 | 601123 |
| Huilliche | Argentina_MitrePeninsula_400BP | Yoruba | 0.275981 | 0.002036 | 135.574 | 823593 |
| Huilliche | Argentina_LagunaToro_2400BP | Yoruba | 0.273181 | 0.002503 | 109.153 | 169287 |
| Huilliche | Chile_WesternArchipelago_800BP | Yoruba | 0.271444 | 0.00192 | 141.343 | 1389101 |
| Huilliche | Peru_Chinchawas_1000BP_MH | Yoruba | 0.271207 | 0.002332 | 116.281 | 344965 |
| Huilliche | Chile_PicaOcho_700BP | Yoruba | 0.270511 | 0.002276 | 118.835 | 458407 |
| Huilliche | Chile_Pukara_700BP_LIP | Yoruba | 0.270369 | 0.002392 | 113.041 | 291363 |
| Huilliche | Peru_Paracas_2250BP_EH | Yoruba | 0.270326 | 0.005571 | 48.523 | 13311 |
| Huilliche | Argentina_NorthTierradelFiego_100BP | Yoruba | 0.270172 | 0.002071 | 130.47 | 1123018 |
| Huilliche | Argentina_LagunaChica_1600BP | Yoruba | 0.270162 | 0.002292 | 117.861 | 550154 |
| Huilliche | Argentina_ArroyoSeco2_7700BP | Yoruba | 0.270074 | 0.00204 | 132.361 | 873735 |
| Huilliche | Peru_Lima_1050BP_MH | Yoruba | 0.269958 | 0.002671 | 101.085 | 88533 |
| Huilliche | Peru_LaGalgada_4100BP | Yoruba | 0.269717 | 0.002151 | 125.371 | 823543 |
| Huilliche | Argentina_BeagleChannel_100BP | Yoruba | 0.269709 | 0.001929 | 139.824 | 976636 |
| Huilliche | Peru_Chinchawas_700BP_LIP | Yoruba | 0.269706 | 0.00207 | 130.316 | 780626 |
| Huilliche | Bolivia_MH_Tiwanaku | Yoruba | 0.2694 | 0.002186 | 123.238 | 337403 |
| Huilliche | Chile_BeagleChannel_800BP | Yoruba | 0.269144 | 0.001994 | 134.989 | 1389293 |
| Huilliche | Peru_Cusco_Torontoy_B_450BP_LH | Yoruba | 0.269118 | 0.002232 | 120.567 | 718205 |
| Huilliche | Bolivia_Iroco_1050BP_MH | Yoruba | 0.269098 | 0.002313 | 116.358 | 539919 |
| Huilliche | Peru_ElBrujo_1300BP_EIP | Yoruba | 0.268969 | 0.002104 | 127.825 | 626441 |
| Huilliche | Peru_Campanayuq_1000BP_MH | Yoruba | 0.268967 | 0.00228 | 117.948 | 227321 |
| Huilliche | Peru_Mesayocpata_450BP_LH | Yoruba | 0.268724 | 0.002562 | 104.888 | 177863 |
| Huilliche | Peru_SanSebastian_600BP_LIP | Yoruba | 0.268397 | 0.002164 | 124.012 | 633877 |
| Huilliche | Peru_SanSebastian_500BP_LH | Yoruba | 0.268373 | 0.002355 | 113.95 | 358746 |
| Huilliche | Peru_Ullujaya_950BP_MH | Yoruba | 0.268329 | 0.002322 | 115.546 | 515038 |

| | | | | | | |
|---|---|---|---|---|---|---|
| Huilliche | Argentina_NorthTierradelFuego_LaArcillosa2_5800BP | Yoruba | 0.268237 | 0.002403 | 111.604 | 630706 |
| Huilliche | Peru_Cuncaicha_4200BP | Yoruba | 0.26811 | 0.002272 | 117.985 | 798226 |
| Huilliche | Peru_Laramate_650BP_LIP | Yoruba | 0.2679 | 0.002044 | 131.041 | 951796 |
| Huilliche | Peru_Ullujaya_1350BP_EIP | Yoruba | 0.267816 | 0.00216 | 123.995 | 512138 |
| Huilliche | Bolivia_Miraflores_1100BP_MH | Yoruba | 0.267754 | 0.002015 | 132.892 | 886713 |
| Huilliche | Peru_Laramate_900BP | Yoruba | 0.267693 | 0.001945 | 137.612 | 894345 |
| Huilliche | Chile_CaletaHuelen_1100BP_MH | Yoruba | 0.267687 | 0.002113 | 126.687 | 476743 |
| Huilliche | Peru_Lima_LIP_650BP | Yoruba | 0.267668 | 0.002092 | 127.979 | 892003 |
| Huilliche | Peru_Lima_EIP_1450BP | Yoruba | 0.267663 | 0.002233 | 119.889 | 321345 |
| Huilliche | Peru_Charrangochayoc_700BP_LIP | Yoruba | 0.267663 | 0.002141 | 124.991 | 728296 |
| Huilliche | Peru_Palpa_950BP_MH | Yoruba | 0.267523 | 0.003566 | 75.028 | 38066 |
| Huilliche | Peru_Cusco_Torontoy_A_450BP_LH | Yoruba | 0.267438 | 0.002145 | 124.686 | 802193 |
| Huilliche | Peru_ElBrujo_850BP_MH-LIP | Yoruba | 0.267097 | 0.002073 | 128.818 | 827395 |
| Huilliche | Peru_Palpa_550BP_LIP | Yoruba | 0.266201 | 0.002044 | 130.227 | 652488 |
| Huilliche | Chile_StraitOfMagellan_100BP | Yoruba | 0.266168 | 0.001961 | 135.763 | 654621 |
| Huilliche | Peru_La_Galgada_600BP_LIP | Yoruba | 0.265979 | 0.005417 | 49.104 | 12201 |
| Huilliche | Peru_Lauricocha_5800BP | Yoruba | 0.265779 | 0.002227 | 119.319 | 581269 |
| Huilliche | Argentina_LagunaChica_6800BP | Yoruba | 0.265463 | 0.002226 | 119.28 | 413848 |
| Huilliche | Peru_Cuncaicha_9000BP | Yoruba | 0.265167 | 0.002295 | 115.535 | 808985 |
| Huilliche | Peru_Totocachi_Tiwanaku | Yoruba | 0.26498 | 0.002168 | 122.244 | 273459 |
| Huilliche | Chile_LosRieles_12000BP | Yoruba | 0.2646 | 0.002441 | 108.4 | 722763 |
| Huilliche | Peru_Cuncaicha_3300BP | Yoruba | 0.264502 | 0.002111 | 125.301 | 707428 |
| Huilliche | Chile_WesternArchipelago_Ayayema_4700BP | Yoruba | 0.26419 | 0.002078 | 127.12 | 1387359 |
| Huilliche | Peru_Lauricocha_3500BP | Yoruba | 0.263909 | 0.002369 | 111.407 | 562112 |
| Huilliche | Peru_RioUncallane_1600BP | Yoruba | 0.263832 | 0.001902 | 138.736 | 1391116 |
| Huilliche | Peru_Lauricocha_8600BP | Yoruba | 0.263764 | 0.002096 | 125.829 | 903895 |
| Huilliche | Peru_Chincha_LH | Yoruba | 0.263055 | 0.001852 | 142.039 | 1390261 |
| Huilliche | Peru_HuacaPrieta_4500BP_LA | Yoruba | 0.262517 | 0.004858 | 54.04 | 18083 |
| Huilliche | Peru_Antaura_Tiwanaku | Yoruba | 0.262378 | 0.00199 | 131.877 | 958683 |
| Huilliche | Bolivia_Akapana_Tiwanaku | Yoruba | 0.261777 | 0.001899 | 137.827 | 954542 |
| Huilliche | Peru_Maucallacta_Tiwanaku | Yoruba | 0.261536 | 0.002074 | 126.115 | 622200 |
| Huilliche | Bolivia_Lukurmata_Tiwanaku | Yoruba | 0.260248 | 0.001857 | 140.157 | 1242298 |
| Huilliche | Bolivia_Pumapunku_Tiwanaku | Yoruba | 0.259898 | 0.002077 | 125.133 | 361554 |

| | | | | | | |
|---|---|---|---|---|---|---|
| Huilliche | Chile_PuntaSantaAna_7300BP | Yoruba | 0.259559 | 0.002115 | 122.745 | 1135309 |
| Huilliche | Peru_Culcunche_Tiwanaku | Yoruba | 0.259209 | 0.002016 | 128.602 | 639369 |
| Huilliche | Peru_Kaillachuro_Unknown | Yoruba | 0.257508 | 0.002096 | 122.872 | 431981 |
| Huilliche | Bolivia_Putuni_Tiwanaku | Yoruba | 0.257489 | 0.002013 | 127.911 | 702037 |
| Huilliche | Bolivia_Monolito_Descabezado_Tiwanaku | Yoruba | 0.256035 | 0.001913 | 133.816 | 945367 |
| Huilliche | Peru_SoroMikayaPatjxa_6800BP | Yoruba | 0.235055 | 0.002568 | 91.516 | 96407 |
| Paran | Peru_Paracas_2250BP_EH | Yoruba | 0.281686 | 0.006071 | 46.398 | 13167 |
| Paran | Peru_Chinchawas_1000BP_MH | Yoruba | 0.281061 | 0.002745 | 102.397 | 341580 |
| Paran | Peru_Charrangochayoc_700BP_LIP | Yoruba | 0.281013 | 0.00241 | 116.621 | 721212 |
| Paran | Peru_Lima_1050BP_MH | Yoruba | 0.280227 | 0.003096 | 90.505 | 87646 |
| Paran | Peru_Chinchawas_700BP_LIP | Yoruba | 0.279925 | 0.002309 | 121.21 | 773765 |
| Paran | Chile_Pukara_700BP_LIP | Yoruba | 0.279403 | 0.00267 | 104.653 | 288563 |
| Paran | Peru_LaGalgada_4100BP | Yoruba | 0.279264 | 0.002501 | 111.679 | 815554 |
| Paran | Peru_Mesayocpata_450BP_LH | Yoruba | 0.279069 | 0.00293 | 95.251 | 176082 |
| Paran | Peru_Campanayuq_1000BP_MH | Yoruba | 0.278986 | 0.002578 | 108.202 | 225053 |
| Paran | Peru_Totocachi_Tiwanaku | Yoruba | 0.277535 | 0.002489 | 111.484 | 268567 |
| Paran | Peru_Laramate_900BP | Yoruba | 0.277486 | 0.002101 | 132.077 | 887939 |
| Paran | Peru_Palpa_950BP_MH | Yoruba | 0.27723 | 0.004075 | 68.037 | 37707 |
| Paran | Peru_Laramate_650BP_LIP | Yoruba | 0.277006 | 0.002217 | 124.931 | 943981 |
| Paran | Peru_Lauricocha_5800BP | Yoruba | 0.276803 | 0.00267 | 103.683 | 575712 |
| Paran | Peru_HuacaPrieta_4500BP_LA | Yoruba | 0.276603 | 0.005722 | 48.343 | 17904 |
| Paran | Peru_Ullujaya_950BP_MH | Yoruba | 0.276068 | 0.00258 | 106.986 | 509895 |
| Paran | Bolivia_MH_Tiwanaku | Yoruba | 0.275927 | 0.002422 | 113.921 | 334093 |
| Paran | Peru_Lima_EIP_1450BP | Yoruba | 0.275729 | 0.002588 | 106.543 | 318219 |
| Paran | Peru_Ullujaya_1350BP_EIP | Yoruba | 0.27559 | 0.002359 | 116.846 | 507254 |
| Paran | Peru_SanSebastian_500BP_LH | Yoruba | 0.275412 | 0.002677 | 102.864 | 355158 |
| Paran | Peru_ElBrujo_1300BP_EIP | Yoruba | 0.275268 | 0.002246 | 122.552 | 620830 |
| Paran | Peru_Cusco_Torontoy_A_450BP_LH | Yoruba | 0.275221 | 0.002271 | 121.197 | 794950 |
| Paran | Peru_SanSebastian_600BP_LIP | Yoruba | 0.275024 | 0.002493 | 110.325 | 627647 |
| Paran | Peru_Lima_LIP_650BP | Yoruba | 0.274979 | 0.002262 | 121.545 | 884507 |
| Paran | Peru_Palpa_550BP_LIP | Yoruba | 0.27422 | 0.002246 | 122.096 | 646502 |
| Paran | Chile_PicaOcho_700BP | Yoruba | 0.273927 | 0.002493 | 109.883 | 453851 |
| Paran | Peru_Lauricocha_8600BP | Yoruba | 0.272739 | 0.002262 | 120.568 | 896057 |

| | | | | | | |
|---|---|---|---|---|---|---|
| Paran | Peru_Cusco_Torontoy_B_450BP_LH | Yoruba | 0.272704 | 0.002535 | 107.566 | 711417 |
| Paran | Peru_ElBrujo_850BP_MH-LIP | Yoruba | 0.272064 | 0.002208 | 123.193 | 820478 |
| Paran | Peru_Antaura_Tiwanaku | Yoruba | 0.271953 | 0.002261 | 120.292 | 941912 |
| Paran | Peru_Lauricocha_3500BP | Yoruba | 0.271736 | 0.002669 | 101.825 | 556906 |
| Paran | Peru_La_Galgada_600BP_LIP | Yoruba | 0.271219 | 0.005964 | 45.473 | 12086 |
| Paran | Peru_Chincha_LH | Yoruba | 0.271118 | 0.002003 | 135.329 | 1373736 |
| Paran | Bolivia_Iroco_1050BP_MH | Yoruba | 0.27096 | 0.002583 | 104.888 | 534688 |
| Paran | Peru_Cuncaicha_3300BP | Yoruba | 0.270874 | 0.002454 | 110.367 | 700602 |
| Paran | Bolivia_Miraflores_1100BP_MH | Yoruba | 0.270786 | 0.002188 | 123.737 | 879402 |
| Paran | Peru_Cuncaicha_4200BP | Yoruba | 0.270453 | 0.002646 | 102.218 | 790685 |
| Paran | Chile_SouthContinent_CerroJohnny_400BP | Yoruba | 0.269844 | 0.002591 | 104.165 | 445522 |
| Paran | Chile_CaletaHuelen_1100BP_MH | Yoruba | 0.269791 | 0.002337 | 115.464 | 472250 |
| Paran | Peru_Cuncaicha_9000BP | Yoruba | 0.269229 | 0.0026 | 103.557 | 801317 |
| Paran | Peru_Maucallacta_Tiwanaku | Yoruba | 0.26892 | 0.002256 | 119.176 | 611445 |
| Paran | Argentina_NorthTierradelFuego_LaArcillosa2_5800BP | Yoruba | 0.268655 | 0.002647 | 101.502 | 624533 |
| Paran | Chile_Conchali_700BP | Yoruba | 0.268512 | 0.002277 | 117.938 | 816033 |
| Paran | Argentina_NorthTierradelFuego_500BP | Yoruba | 0.268115 | 0.002277 | 117.768 | 845393 |
| Paran | Chile_NorthTierradelFuego_100BP | Yoruba | 0.26804 | 0.002628 | 101.996 | 595384 |
| Paran | Argentina_LagunaChica_1600BP | Yoruba | 0.267921 | 0.002489 | 107.636 | 544809 |
| Paran | Peru_Kaillachuro_Unknown | Yoruba | 0.267522 | 0.002271 | 117.806 | 423980 |
| Paran | Peru_RioUncallane_1600BP | Yoruba | 0.26721 | 0.002076 | 128.722 | 1373480 |
| Paran | Chile_LosRieles_5100BP | Yoruba | 0.267139 | 0.002561 | 104.3 | 698439 |
| Paran | Argentina_MitrePeninsula_400BP | Yoruba | 0.267062 | 0.002285 | 116.901 | 817125 |
| Paran | Argentina_LagunaToro_2400BP | Yoruba | 0.267033 | 0.002803 | 95.281 | 167526 |
| Paran | Peru_Culcunche_Tiwanaku | Yoruba | 0.267017 | 0.002295 | 116.351 | 628160 |
| Paran | Bolivia_Akapana_Tiwanaku | Yoruba | 0.266993 | 0.002136 | 124.976 | 938955 |
| Paran | Argentina_BeagleChannel_1900-500BP | Yoruba | 0.266707 | 0.002279 | 117.046 | 864737 |
| Paran | Chile_LosRieles_12000BP | Yoruba | 0.266208 | 0.002701 | 98.573 | 715821 |
| Paran | Bolivia_Lukurmata_Tiwanaku | Yoruba | 0.26387 | 0.002065 | 127.793 | 1223453 |
| Paran | Argentina_ArroyoSeco2_7700BP | Yoruba | 0.263542 | 0.002176 | 121.106 | 866838 |
| Paran | Bolivia_Pumapunku_Tiwanaku | Yoruba | 0.262931 | 0.002413 | 108.968 | 355351 |
| Paran | Argentina_NorthTierradelFiego_100BP | Yoruba | 0.262292 | 0.002303 | 113.907 | 1103714 |
| Paran | Argentina_LagunaChica_6800BP | Yoruba | 0.262258 | 0.002419 | 108.399 | 409981 |

| | | | | | | |
|---|---|---|---|---|---|---|
| Paran | Chile_WesternArchipelago_800BP | Yoruba | 0.261991 | 0.002179 | 120.211 | 1367644 |
| Paran | Argentina_BeagleChannel_100BP | Yoruba | 0.261102 | 0.002201 | 118.647 | 961808 |
| Paran | Bolivia_Putuni_Tiwanaku | Yoruba | 0.26083 | 0.002263 | 115.241 | 689741 |
| Paran | Chile_WesternArchipelago_Ayayema_4700BP | Yoruba | 0.259668 | 0.002302 | 112.782 | 1362892 |
| Paran | Chile_BeagleChannel_800BP | Yoruba | 0.259279 | 0.002215 | 117.069 | 1367528 |
| Paran | Chile_PuntaSantaAna_7300BP | Yoruba | 0.258178 | 0.002327 | 110.958 | 1115056 |
| Paran | Bolivia_Monolito_Descabezado_Tiwanaku | Yoruba | 0.257546 | 0.002185 | 117.849 | 929091 |
| Paran | Chile_StraitOfMagellan_100BP | Yoruba | 0.255185 | 0.002175 | 117.331 | 643664 |
| Paran | Peru_SoroMikayaPatjxa_6800BP | Yoruba | 0.237705 | 0.002828 | 84.051 | 94537 |

**Table S6. Summary of DATES results for the El Plomo individual.**

| SOURCE1 | N | SOURCE2 | N | MEAN | STD_ERROR | Z | NRMDS | YEARS | YEARS BCE/CE |
|---|---|---|---|---|---|---|---|---|---|
| Huilliche | 17 | Paran | 2 | 260.702 | 83.146 | 3.135 | 0.305 | | |
| Huilliche | 17 | Cusco | 3 | 49.546 | 18.745 | 2.643 | 0.634 | 1387.3 | 72.7 CE |
| Huilliche | 17 | Cusco2 | 7 | 907.358 | 430.674 | 2.107 | 0.433 | | |
| Huilliche | 17 | Aymara | 40 | 145.276 | 42.839 | 3.391 | 0.45 | 4067.7 | 2607.7 BCE |
| Chile_LosRieles_5100BP | 1 | SouthPeruHighlands | 14 | 125.658 | 44.985 | 2.793 | 0.656 | 3518.4 | 2058.4 BCE |
| Chile_LosRieles_5100BP | 1 | NorthPeruHighlands | 5 | 818.985 | 707.274 | 1.158 | 0.435 | | |
| Chile_Conchali_700BP | 2 | SouthPeruHighlands | 14 | 135911.093 | 30600.695 | 4.441 | 0.945 | | |
| Chile_Conchali_700BP | 2 | NorthPeruHighlands | 5 | 556.469 | 530.537 | 1.049 | 0.319 | | |

* Best models highlighted in yellow

**Table S7. D-statistics of the form D(Huilliche, Chile_Conchali_700BP; X; Yoruba).** X represents one present-day populations from central-south and coastal Peru.

| H1 | H2 | H3 | H4 | D-stat | SE | Z-score | BABA | ABBA | SNPs |
|---|---|---|---|---|---|---|---|---|---|
| Huilliche | Chile_Conchali_700BP | Paran | Yoruba | 0.0023 | 0.003215 | 0.717 | 40695 | 40507 | 866482 |
| Huilliche | Chile_Conchali_700BP | Cusco | Yoruba | 0.0015 | 0.002953 | 0.51 | 40693 | 40570 | 866575 |
| Huilliche | Chile_Conchali_700BP | Cusco2 | Yoruba | 0.0018 | 0.002633 | 0.683 | 40658 | 40512 | 866575 |
| Huilliche | Chile_Conchali_700BP | Aymara | Yoruba | 0.0015 | 0.002367 | 0.614 | 40624 | 40506 | 866575 |
| Huilliche | Chile_Conchali_700BP | Eten | Yoruba | 0.0019 | 0.00272 | 0.706 | 40649 | 40493 | 866575 |
